# DPHL: A pan-human protein mass spectrometry library for robust biomarker discovery

**DOI:** 10.1101/2020.02.03.931329

**Authors:** Tiansheng Zhu, Yi Zhu, Yue Xuan, Huanhuan Gao, Xue Cai, Sander R. Piersma, Thang V. Pham, Tim Schelfhorst, Richard R Goeij De Haas, Irene V. Bijnsdorp, Rui Sun, Liang Yue, Guan Ruan, Qiushi Zhang, Mo Hu, Yue Zhou, Winan J. Van Houdt, T.Y.S Lelarge, J. Cloos, Anna Wojtuszkiewicz, Danijela Koppers-Lalic, Franziska Böttger, Chantal Scheepbouwer, R.H Brakenhoff, G.J.L.H. van Leenders, Jan N.M. Ijzermans, J.W.M. Martens, R.D.M. Steenbergen, N.C. Grieken, Sathiyamoorthy Selvarajan, Sangeeta Mantoo, Sze Sing Lee, Serene Jie Yi Yeow, Syed Muhammad Fahmy Alkaff, Nan Xiang, Yaoting Sun, Xiao Yi, Shaozheng Dai, Wei Liu, Tian Lu, Zhicheng Wu, Xiao Liang, Man Wang, Yingkuan Shao, Xi Zheng, Kailun Xu, Qin Yang, Yifan Meng, Cong Lu, Jiang Zhu, Jin’e Zheng, Bo Wang, Sai Lou, Yibei Dai, Chao Xu, Chenhuan Yu, Huazhong Ying, Tony Kiat-hon Lim, Jianmin Wu, Xiaofei Gao, Zhongzhi Luan, Xiaodong Teng, Peng Wu, Shi’ang Huang, Zhihua Tao, N. Gopalakrishna Iyer, Shuigeng Zhou, Wenguang Shao, Henry Lam, Ding Ma, Jiafu Ji, Oi Lian Kon, Shu Zheng, Ruedi Aebersold, Connie R. Jimenez, Tiannan Guo

## Abstract

To answer the increasing need for detecting and validating protein biomarkers in clinical specimens, proteomic techniques are required that support the fast, reproducible and quantitative analysis of large clinical sample cohorts. Targeted mass spectrometry techniques, specifically SRM, PRM and the massively parallel SWATH/DIA technique have emerged as a powerful method for biomarker research. For optimal performance, they require prior knowledge about the fragment ion spectra of targeted peptides. In this report, we describe a mass spectrometric (MS) pipeline and spectral resource to support data-independent acquisition (DIA) and parallel reaction monitoring (PRM) based biomarker studies. To build the spectral resource we integrated common open-source MS computational tools to assemble an open source computational workflow based on Docker. It was then applied to generate a comprehensive DIA pan-human library (DPHL) from 1,096 data dependent acquisition (DDA) MS raw files, and it comprises 242,476 unique peptide sequences from 14,782 protein groups and 10,943 SwissProt-annotated proteins expressed in 16 types of cancer samples. In particular, tissue specimens from patients with prostate cancer, cervical cancer, colorectal cancer, hepatocellular carcinoma, gastric cancer, lung adenocarcinoma, squamous cell lung carcinoma, diseased thyroid, glioblastoma multiforme, sarcoma and diffuse large B-cell lymphoma (DLBCL), as well as plasma samples from a range of hematologic malignancies were collected from multiple clinics in China, the Netherlands and Singapore and included in the resource. This extensive spectral resource was then applied to a prostate cancer cohort of 17 patients, consisting of 8 patients with prostate cancer (PCa) and 9 with benign prostate hyperplasia (BPH), respectively. Data analysis of DIA data from these samples identified differential expressions of FASN, TPP1 and SPON2 in prostate tumors. Thereafter, PRM validation was applied to a larger PCa cohort of 57 patients and the differential expressions of FASN, TPP1 and SPON2 in prostate tumors were validated. As a second application, the DPHL spectral resource was applied to a patient cohort consisting of samples from 19 DLBCL patients and 18 healthy individuals. Differential expressions of CRP, CD44 and SAA1 between DLBCL cases and healthy controls were detected by DIA-MS and confirmed by PRM. These data demonstrate that the DPHL supported that DIA-PRM MS pipeline enables robust protein biomarker discoveries.

## INTRODUCTION

The recent development of high throughput genomic sequencing techniques, as well as methods for the global expression analysis of biomolecules has enabled identification of a number of oncological biomarkers from clinical samples, and advanced the field of cancer precision medicine [1–4]. Novel diagnostic/prognostic protein markers for colorectal [5, 6], breast [7], ovarian [8] and gastric tumors [9] have been identified through shotgun proteomics [10], and plasma proteomes were reported for 1500 obese patients [11]. Sequential window acquisition of all theoretical fragment ion spectra mass spectrometry (SWATH-MS) is a data independent acquisition technique that combines the multiplexing ability of shotgun proteomics with the high-precision data analysis of selected reaction monitoring (SRM), and can quantify proteomes using single-shot MS/MS analysis [12, 13]. The SWATH/DIA data sets are analyzed through spectral libraries using software tools like OpenSWATH [14, 15], DIA-Umpire [16], Group-DIA [17], Skyline [18], Spectronaut [19]. Most of these tools generate comparable results [15] and requires *a prior* spectral libraries. A pan-human spectral library (PHL) that was designed to aid in SWATH data processing has been developed to analyze SWATH maps generated by TripleTOF MS [20] by using open-source computational programs [1, 14], then the error rates of peptide and protein identification in large-scale DIA analyses has been statistically controlled [21]. The development of these tools has extended the application of SWATH-MS to diverse clinical samples including plasma [22], and the prostate [23] and liver [24] cancer tissues.

Despite these advances, the implementation of DIA-MS on widely used Orbitrap instruments is currently limited due to the lack of non-commercial tools to build spectral libraries. Theoretically one could build a spectral library based on the established protocol for TripleTOF data [1], however in practice an optimal and robust pipeline for Orbitrap data is missing, as we have implemented in this work. Further, it has been demonstrated that the library from TripleTOF led to fewer protein identifications than that from Orbitrap [25]. Moreover, there is no bioinformatics pipeline to couple DIA-MS and PRM-MS for validation, and a comprehensive human spectral library resource for Orbitrap data is yet to be established. Spectronaut has been developed to support the generation of DIA spectral libraries and analysis of DIA data sets against these libraries [19], however, it is only commercially available. Parallel computing is only available for OpenSWATH software tools till now. To extend the application of large-scale DIA-MS on Orbitrap instruments, an open-source workflow is in great need to build a pan-human spectral library for DIA files generated for cancer biomarker discovery. Further, the open-source workflow and the spectral library are essential to validate the candidate protein biomarkers by PRM that is a more recently developed technique with higher sensitivity and specificity than SWATH/ DIA, albeit with limited throughput [26].

Here, we developed an open-source computational pipeline to build spectral libraries from Orbitrap spectral data, and generated a comprehensive DIA Pan-Human Library (DPHL) from 16 different human cancer types. In addition, we have also provided a Docker resource to integrate this pipeline to the data-dependent acquisition (DDA) spectral scans, which allows an easy and automatic expansion of the library by incorporating more MS data generated from ongoing studies. Finally, to validate its applicability in DIA and PRM, we applied the DPHL to identify differentially expressed proteins in the samples from a prostate cancer and a DLBCL cohort.

## RESULTS AND DISCUSSION

### Shotgun proteomics data of tumor tissues and plasma samples

To build a DIA spectral library for Orbitrap data which can also be used for PRM assay generation, we obtained shotgun proteomics data from two laboratories in China and the Netherlands that use Q Exactive HF mass spectrometers and consistent experimental conditions (see Materials and Methods section). A total of 1,096 raw MS data files were collected from a range of samples that included tissue biopsies from prostate cancer, cervical cancer, colorectal cancer, hepatocellular carcinoma, gastric cancer, lung adenocarcinoma, squamous cell lung carcinoma, thyroid diseases, glioblastoma multiforme, sarcoma and DLBCL. Further, blood plasma samples from acute myelocytic leukemia (AML), acute lymphoblastic leukemia (ALL), chronic myelogenous leukemia (CML), multiple myeloma (MM), myelodysplastic syndrome (MDS) and DLBCL patients, and the human chronic myelogenous leukemia cell line K562 were also analyzed and the data were included in the library. The sample types and their DDA files are summarized in Figure 1A and Supplementary Table S1A. Comparison of DDA files acquired from the Guo lab and the Jimenez lab is provided in Supplementary Note 1.

**Figure 1.**
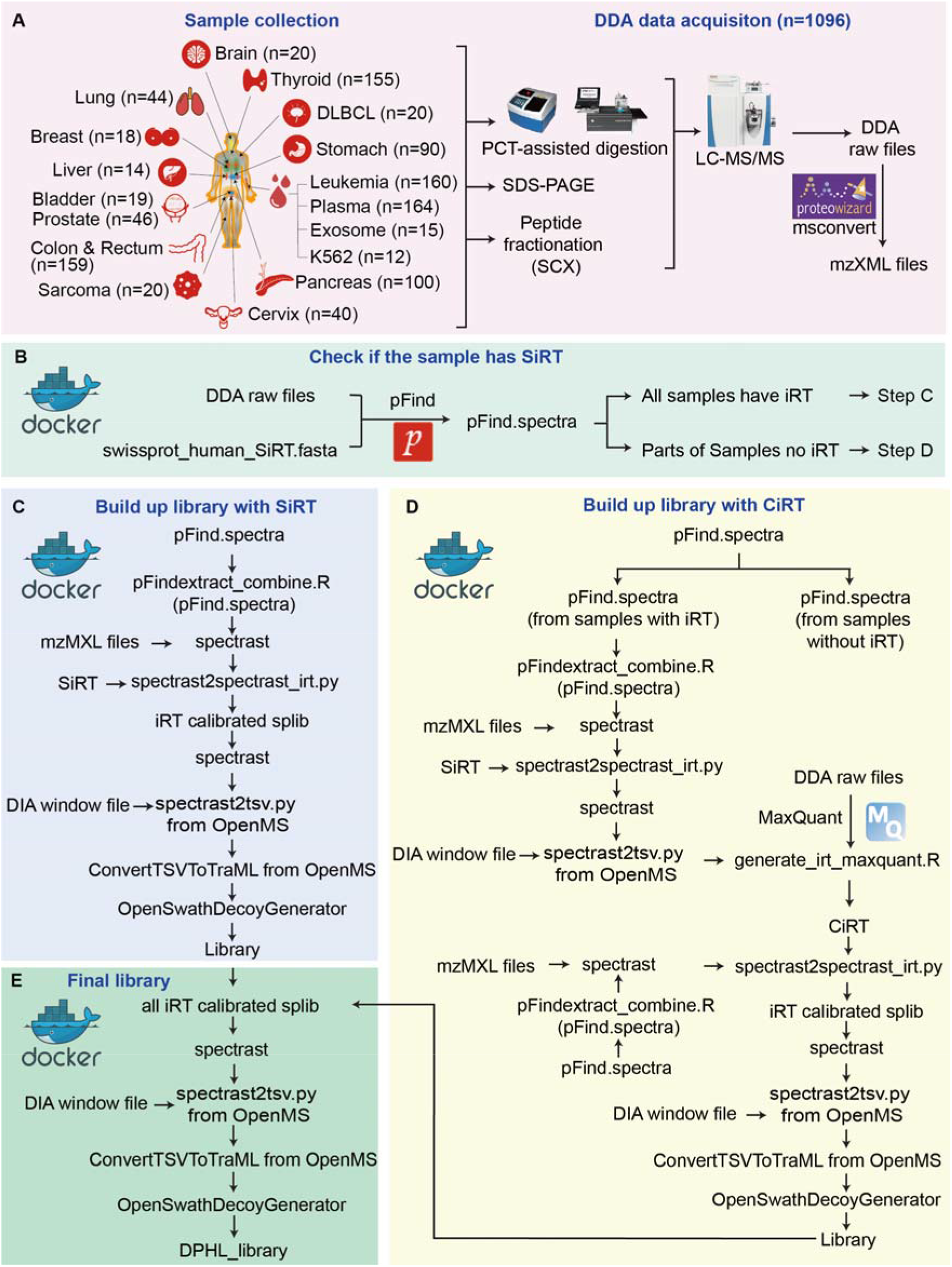
Workflow for building the DPHL. (A) Schematic representation of DDA shotgun proteomics data acquisition. Numbers in parentheses indicate the number of DDA files per tissue type. B-E. Computational pipeline for building DIA spectral library. (B) Protein identification and iRT detection from DDA raw files using pFind. (C) SiRT detection and calibration. (D) CiRT detection and calibration. (E) Generation of the DPHL library. Details of the commands are presented in Supplementary Note 1.

### Open-source computational pipeline for building DIA/PRM spectral library

The conventional OpenMS and OpenSWATH pipeline [14] requires sophisticated installation which relies on multiple existing software packages. A Docker image largely facilitate the installation process. We developed an open-source Docker image with all the pre-installed pipelines and its dependent packages to democratize the generation of DIA/PRM spectral libraries. The workflow of this computational pipeline is shown in Figure 1B. Briefly, the DDA files were first centroided and converted to mzXML using MSconvert from ProteoWizard [27], and pFind [28] was used to identify the relevant peptides and proteins in the protein database. The shotgun data from each tissue type was processed separately. We wrote two scripts – pFindextract.R and addRT.py – to extract the retention time (RT), peptide sequence, charge state, protein name and identification score for each peptide precursor. Spectrast version 5.0 [29] was used to generate consensus spectra of peptides for each tissue type to build the library, spectrast2spectrast_irt.py [30] was used for RT calibration, and spectrast2tsv.py [14] for selecting the top six fragments for each peptide precursor. Decoy assays were generated using OpenSwathDecoyGenerator from OpenSWATH software [14].

For both, library building and SWATH/DIA analysis, the peptide samples were usually spiked with a synthetic iRT peptides mixture (SiRT) [31] to calibrate the retention time, and the SWATH library building workflow [1] was also applied to these samples. For the samples without SiRT spike-in, we employed tools to identify the conserved high-abundance peptides with common internal retention time (CiRT) [30]. The peptides of each tissue type had to fulfill the following criteria to be considered as CiRTs peptides: (1) proteotypic, (2) amino acid sequences with no modification, (3) signal intensities above the 3rd quartile of all quantified peptide precursors, (4) charge +2 or +3, and (5) uniformly distributed retention time across the entire LC gradient. Following these criteria, we implemented codes dividing the LC gradient window into 20 bins, and selecting one peptide for each bin. Thereby we selected 20 CiRT peptides for each tissue type. The CiRT of the different tissue types are shown in Supplementary Table S2. The TraML format of the CiRT peptides are provided in Supplementary File S1. The CiRT peptides can either be used synergistically with exogenous SiRT standards or as an alternative RT standard in the respective samples. We expect these CiRT peptides to be of wide use in future DIA experiments for these clinical tissue samples.

Since the current version of the pFind software does not support the quantification of identified peptides, CiRT peptides were selected from a representative DDA data set which was analyzed by MaxQuant (version 1.6.2) [32]. We then wrote the generate_CiRT script to analyze the peptides.txt files from the MaxQuant search results, and generated the tissue-specific CiRTs. The latter was used to replace SiRTs in the command spectrast2spectrast_irt.py [30]. For RT calibration, we used the spectrast2spectrast_irt.py converter script on the SiRT or CiRT peptides. Similarly, spectrast was then used to build a consensus library, and spectrast2tsv.py and OpenSwathDecoyGenerator [14] to append decoy assays into the library. The computational pipeline is illustrated and explained in more detail in Supplementary Note 2.

### Build and characterization of the DPHL library

We first characterized the content of the newly-generated DPHL library in terms of the peptide and proteins identifications and compared it to the PHL library for SWATH [20]. The DPHL library includes 359,627 transition groups (peptide precursors), 242,476 unique peptide sequences, 14,782 protein groups, and 10,943 proteotypic SwissProt proteins (Figure 2A). And DPHL contains 2842 protein groups and 1173 proteotpyic SwissProt proteins identified from a single peptide. The two libraries share 9,241 unique proteins, which represent 84.4% of the DPHL and 89.5% of the PHL contents, respectively (Figure 2A). The DPHL library includes more transition groups, unique peptide sequences and protein groups compared to the PHL SWATH library (Figure 2A). Proteins in DPHL are of higher sequence coverage (Supplementary Figure S1), enabling better measurement of specific domains of proteins.

**Figure 2.**
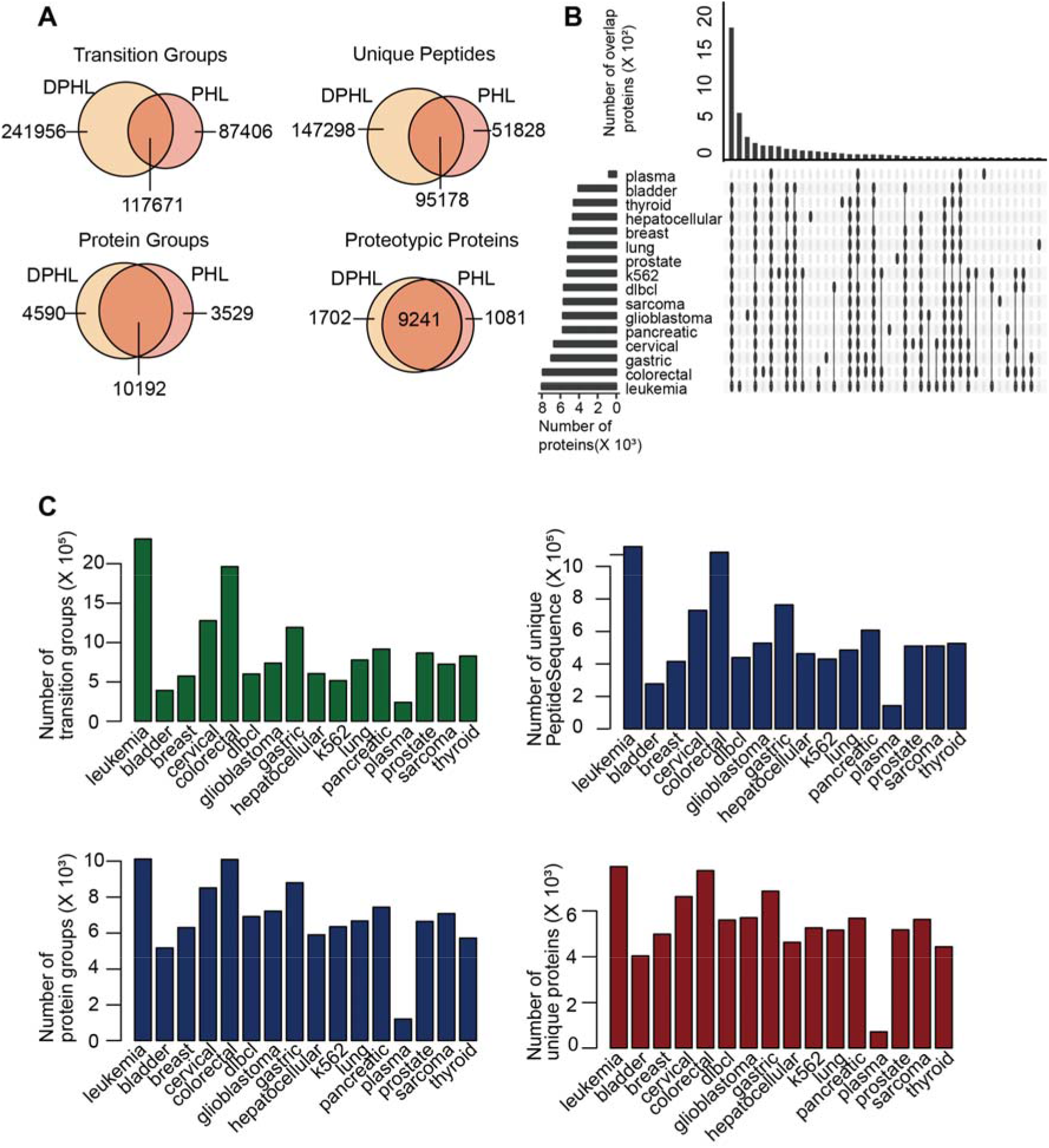
Comparison of DPHL and PHL. (A) Venn diagram showing the comparison of transition groups, unique peptide sequences, protein groups, and proteins in DPHL and PHL. (B) Visualization of set intersections using R package UpSet. (C) The bar plots display the number of transition groups (peptide precursors), unique peptide sequences, protein groups, proteotypic SwissProt proteins in DPHL library for each sample type.

We then counted the number of peptide precursors, unique peptide sequences, and protein groups for each of the 16 sample types (Figure 2B) and found that the solid tissues, but not the plasma samples, shared a large number of proteins. The leukemia samples had the highest number of peptides and proteins due to the higher number of DDA files (n = 160) available. The plasma samples had, as expected, the lowest number of peptides and proteins due to the dominance of high abundance proteins. Cumulative plots of peptides and proteins of the 16 types of cancer (tissue, plasma and cell line) are shown in Supplementary Figure S1a and Supplementary Figure S1b. There was a significant increase in the number of transition groups when DDA data was added from different tissue types (Supplementary Figure S2A), while the increase in the number of proteins was relatively less (Supplementary Figure S2B). We further investigated the increase of peptide precursors and proteins in two well sampled tissue type and found that this DPHL library is not yet complete, probably due to semi-tryptic peptides and missed cleavages due to biological heterogeneity (Supplementary Figure S2C, S2D), awaiting for future expansion with more spectral data.

Next, we analyzed the biological content of the DPHL library. To investigate the biological coverage of this DPHL, we did GO (Gene Ontology) enrichment analysis using R package clusterProfiler, as shown in Supplementary Figure S3, demonstrating that our DPHL covers proteins with diverse molecular functions.

The kinases were next characterized using KinMap [33], an online tool that links the biochemical, structural and disease association data of individual kinases to the human kinome tree. A total of 340 kinases (63.2% out of 538 known protein kinases) identified in DPHL were plotted in the KinMap tree. As shown in Supplementary Figure S4, DPHL covers all the major branches of the kinome tree. More characteristics of the kinases in DPHL are show in Supplementary Figure S5. Transcription factors (TFs) are special proteins that bind target DNA sequence to regulate and control gene transcription. TFs are extremely important to disease genesis, development and disease progression. We matched our DPHL library to the 1639 TFs from the Human Transcription Factors database [34], and found that the DPHL covers 33.0% of the known TFs (Supplementary Figure S6).

### Application of the DPHL library to prostate cancer tissue samples

Next we apply the DPHL library to analyze representative clinical sample cohorts. First, we procured prostate tissue samples from 17 patients, consisting of 8 prostate cancers (PCa) and 9 cases of benign prostate hyperplasia (BPH) (Supplementary Table S3), and analyzed them by QE-HF MS operated in DIA mode. The peptides were separated on a 60 min LC gradient. Two additional technical replicates were randomly selected for each patient group. Twenty-four DIA files were thus acquired, 4,785 protein groups, 4,391 SwissProt proteins and 3,723 proteotypic proteins were identified from 37,581 peptide precursors that were searched against the DPHL library using the CiRT strategy (Figure 3A). Figure 3B shows that proteins were identified at a high degree of reproducibility across the samples tested. The SiRT and CiRT strategies achieved comparable performance (Figure 3C). T-SNE[35] plots show that PCa and BPH were clearly distinguished by the data analyzed by both, the CiRT and SiRT strategies (Figure 3D).

**Figure 3.**
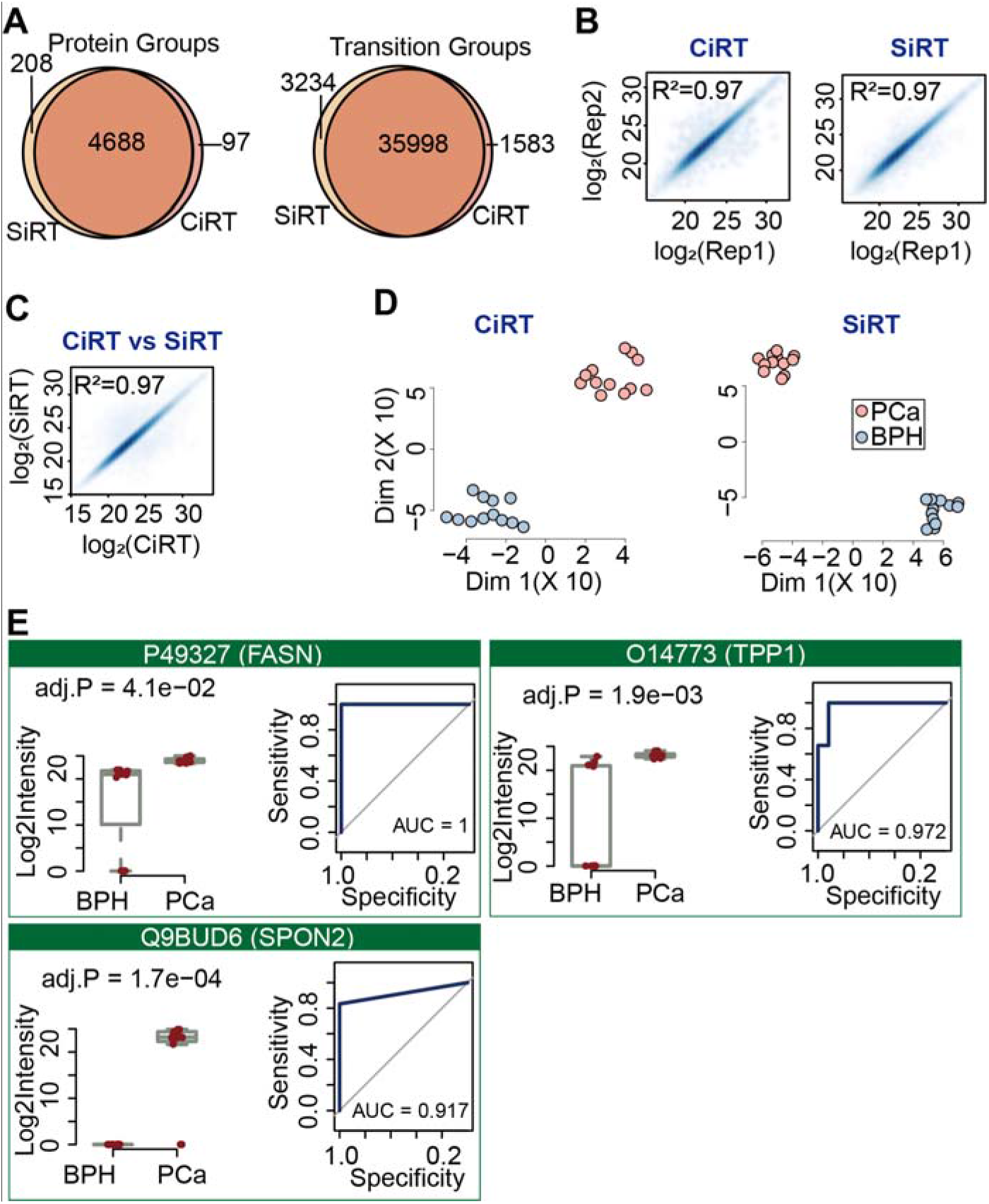
Prostate cancer proteome using 60-min gradient DIA. (A) Peptide and protein identification using SiRT and CiRT. (B) Technical reproducibility of proteome matrix using CiRT and SiRT. (C) Comparison of quantified peptide precursors using the SiRT and CiRT methods. (D) 2D plane t-SNE plot of disease classes, color coded by sample type using CiRT and SiRT. (E) Boxplots and ROC curves showing the significantly dysregulated proteins; p-values are shown under each protein name.

Of the 3,723 identified proteins, 1,555 (1,451 up, 104 down) showed significant differential abundance (Benjamini-Hochberg (BH) adjusted p-values <0.05 and intensity fold-change higher than 2 or lower than 0.5) using CiRT compare to 2,109 (1,954 up and 155 down) proteins using SiRT (see Supplementary Table S3E-S3F). And we used Random forest to select the top 400 most important proteins contributing to the separation of benign and malignant samples, followed by metascape [36] and DAVID [37] for pathway enrichment analysis. We then identified four representative biomarker candidates based on their molecular functions, including fatty acid synthase (P49327, FASN), tripeptidyl-peptidase 1 (O14773, TPP1), and spondin-2 (Q9BUD6, SPON2). FASN, TPP1 and SPON2 were significantly regulated. FASN overexpression has been reported to be associated with poor prognosis in prostate cancers [38]. TPP1 regulates single-stranded telomere DNA binding and telomere recruitment, thus maintaining telomere stability [39–41]. Since genomic instability drives prostate cancer progression from androgen-dependence to castration resistance [42], TPP1 is a promising biomarker [43]. SPON2 is a cell adhesion protein which plays a role in tumor progression and metastasis, and was reported as a serum biomarker [44–46]. The boxplots and ROC curves of these proteins are shown in Figure 3E.

### Application to diffuse large B cell lymphoma (DLBCL) plasma samples

Plasma is widely used in clinical diagnosis for its convenient access. Here we applied the DIA mass spectrometry and the DPHL resource to analyze the plasma samples from DLBCL patients. The plasma samples were procured from 19 DLBCL patients and 18 healthy individuals (Supplementary Table S5). Each unfractionated and un-depleted plasma sample was trypsinized and the resulting peptides were separated on a 20 min LC gradient and measured by DIA-MS on a QE-HF instrument. A total of 7,333 peptide precursors were identified by searching the data against the DPHL plasma subset library using the CiRT strategy with high technical reproducibility (R^2^ = 0.96, Figure 4A). We identified 507 protein groups and 304 proteotypic proteins. More detailed information per sample was show in Supplementary Figure S7. The DLBCL samples were clearly distinguished from the healthy control samples by t-SNE analysis of the quantified proteome (Figure 4B), indicating that our workflow can distinguish DLBCL patients from healthy individuals based on their plasma proteomes.

**Figure 4.**
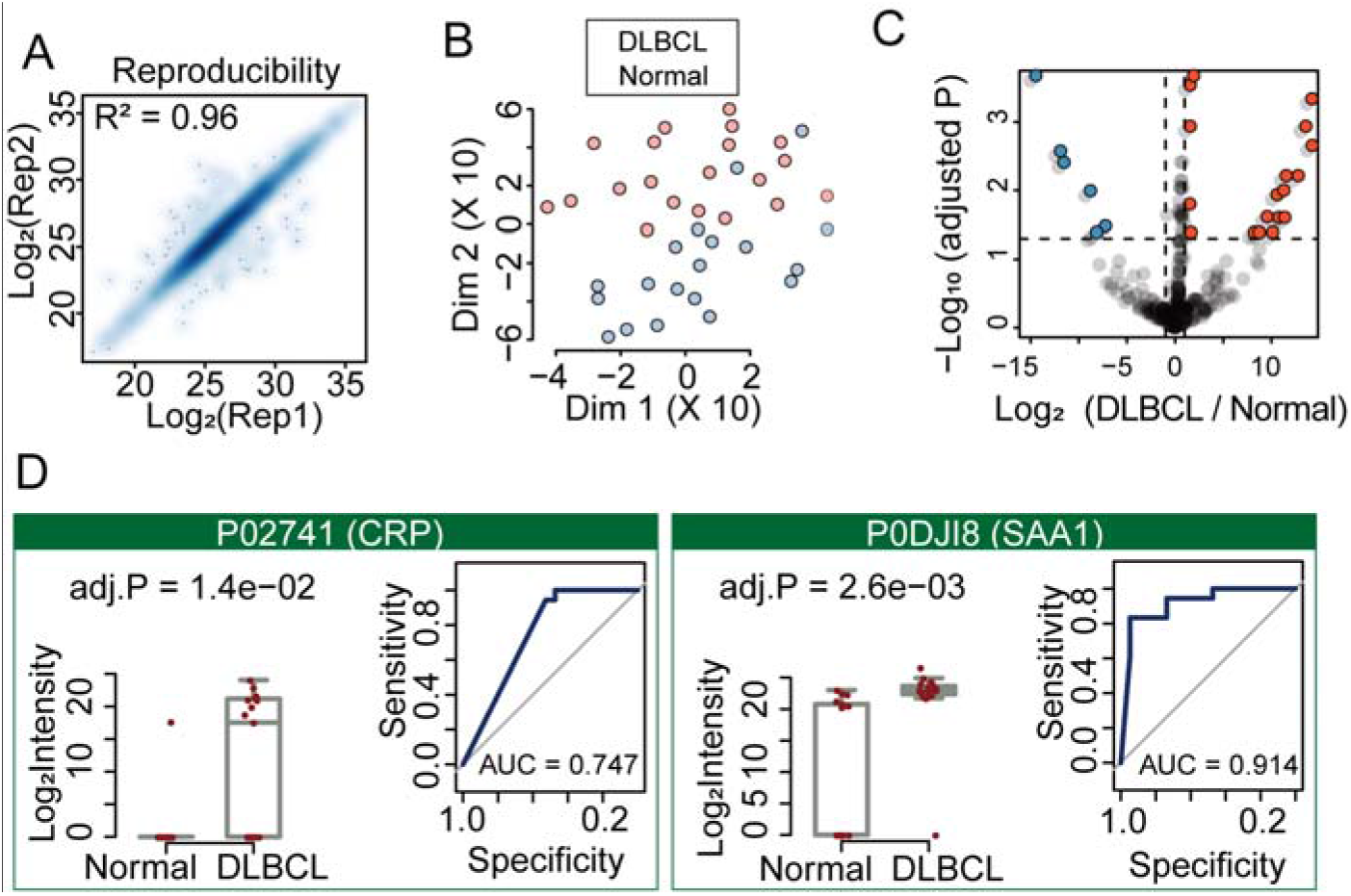
DIA analysis of plasma samples from DLBCL patients and healthy subjects. (A) 2D plane t-SNE plot showing the proteomes are separated. (B) Volcano plot showing significantly down-regulated (blue) and up-regulated (red) proteins in the 37 plasma samples. (C) Technical reproducibility for protein quantification of four plasma samples from DLBCL patients and healthy subjects. (D) Each box shows the expression of a protein biomarker candidate. Left panel: boxplots show the expression difference with P values computed using Student’s *t* test adjusted by the Benjamini-Hochberg method. Right panel: ROC curves of the respective dysregulated protein.

After comparing the DLBCL/healthy (or normal) plasma proteomes using *t*-test with same criteria as the prostate cohort, we identified 24 differential proteins (18 up and 6 down, Supplementary Table S5D), from which we choose three biomarker candidates (Figure 4C) which were closely associated to DLBCL among these 24 proteins based on literature, including C-reactive protein (CRP), CD44 and serum amyloid A1 (SAA-1). CRP is an indicator of the inflammatory response and has prognostic value in various solid tumors, including DLBCL [47]. The hyaluronic acid receptor CD44 and SAA-1 have been previously identified as prognostic biomarkers for DLBCL [48] [49]. The boxplots and ROC curves of these proteins are shown in Figure 4D. Taken together, our workflow can identify potential prognostic biomarkers of DLBCL.

### DPHL-assisted protein validation using PRM

We then validated the candidate biomarkers using PRM, a highly specific and sensitive analytical method that can systematically and precisely quantify well-defined sets of peptides in complex samples. The DPHL spectra were used to develop PRM assays using Skyline [18].

#### Validation in prostate samples

To validate the DIA results of the prostrate samples, we included another independent cohort, thereby increasing the total number of samples to 73 from 57 patients (Supplementary Table S4). The two best flying peptides were selected for each protein to measure the abundance of FASN, TPP1 and SPON2 (Figure 5). As shown in Figure 3E and Figure 5, the PRM well confirmed the DIA results. As a representative example, the peak areas of protein TPP1 (O14773) across all samples are shown in Supplementary Figure S8.

**Figure 5.**
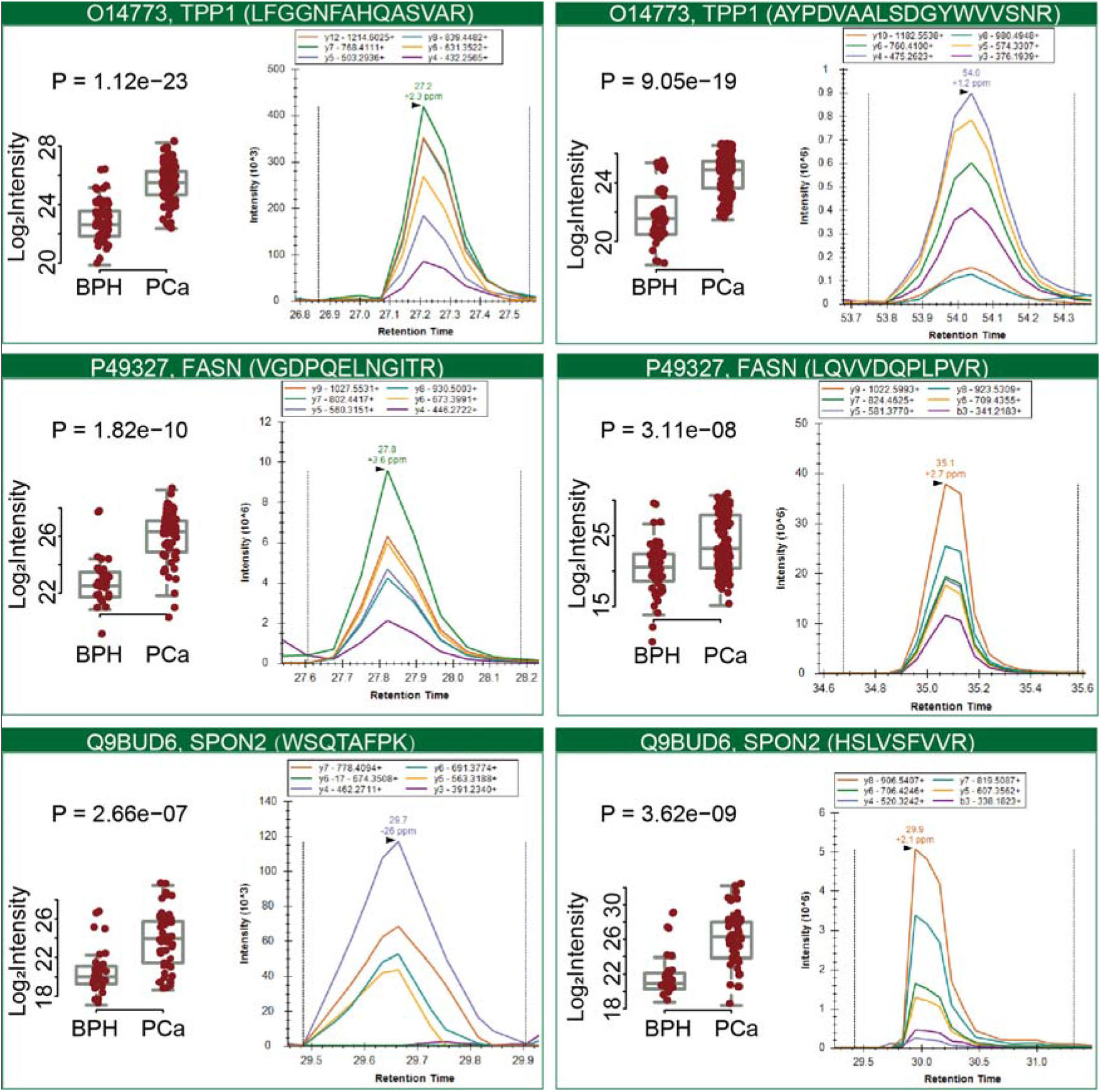
PRM validation of eight peptides in 73 prostate samples. In each box, the left panel shows the log_2_ intensity of eight representative peptides across 73 prostate samples, while the right panel depicts a representative peak group for the respective peptide. P values are computed using Student’s *t* test.

#### Validation in plasma samples

The putative DLBCL biomarkers P02741 (CRP) and P0DJI8 (SAA1) that were identified from the DIA dataset were selected for PRM validation. Skyline was used to visualize characteristic peptides for CRP and SAA1. One of the best flying peptides were selected for CRP and SAA1 to measure the abundance of each protein, respectively (Supplementary Figure S9). The peak groups of the fragment ions were manually curated. As shown in Figure S9, both proteins are highly upregulated in DLBCL patients compared to healthy individuals, confirming the results obtained by DIA (Figure 4D). As an example, the peak areas of peptide ESDTSYVSLK (m/z 564.77) of CRP (P02741) across all samples are shown in Supplementary Figures S10.

## CONCLUSION

In this study, we have developed an open-source platform consisting of a computational pipeline to generate spectral libraries for DIA and PRM analyses on Orbitrap instruments. We also reported a reference spectral library, which can be used to identify and validate protein biomarkers in clinical samples using DIA-MS. With over 370,000 peptide precursors and more than 10,000 proteotypic SwissProt proteins, the DPHL library is the most comprehensive SWATH/DIA library built to date, and allows convenient partitioning into tissue- and disease-specific sub-libraries. Additionally, the DPHL is specifically designed for protein measurement of clinical samples including tissues and plasma, while the PHL is mainly for cell lines and synthetic peptides. Using this approach, we were able to analyze proteomes of 20 human tissue and 40 plasma proteomes per MS instrument per day. We will continue to generate additional DDA files from more types of human tumors with the ambition of incorporating internal and external data to create a comprehensive resource reflecting tumor heterogeneity that enables biomarker discovery as a mission of the Human Proteome Organization Cancer HPP project [50]. By appending these results to the DPHL, we can increase the human proteome coverage. The DPHL is not only applicable to open-source SWATH/DIA analysis tools like OpenSWATH, but also to other tools including Spectronaut and Skyline.

## MATERIALS AND METHODS

All chemicals were from Sigma unless otherwise stated. All HPLC/MS grade reagents for mass spectrometry were from Thermo Fisher.

### Clinical samples

Formalin-fixed paraffin-embedded (FFPE), fresh or fresh frozen (FF) tissue biopsies from prostate cancer, cervical cancer, colorectal cancer, hepatocellular carcinoma, gastric cancer, lung adenocarcinoma, squamous cell lung carcinoma, thyroid diseases, glioblastoma multiforme, sarcoma, and DLBCL were analyzed in this study. Human plasma samples from a range of types of leukemia, lymphoma, plasma cell disorders, anemia, and DLBCL were also included. The human chronic myelocytic leukemia cell line, K562, was present in the dataset. The details about the samples are described in Supplementary Table S1a. Ethics approvals for this study were obtained from the Ethics Committee or Institutional Review Board of each participating institution.

#### Chinese cancer tissue cohorts

Prostate cancer FFPE samples were acquired from the Second Affiliated Hospital of Zhejiang University School of Medicine. The first cohort included 3 PCa patients and 3 patients with BPH was used for DPHL library building. The second cohort containing 8 PCa patients and 9 BPH patients was selected for DIA-MS proteotyping. For each patient, four tissue biopsies (punch 1×1×5 mm^3^) from the same region were procured for the subsequent PCT-SWATH/DIA workflow for targeted quantitative proteomics profiling. Besides the second cohort, a third cohort included 53 patients (16 BPH and 57 PCa) was also included for PRM validation. PRM and DIA analyses were performed in technical duplicate. Information about samples of patient used for DIA and PRM measurements are described in Supplementary Table S3 and Supplementary Table S4.

The colorectal tissue cohort (CRC) was acquired from histologically confirmed tumors at the First Affiliated Hospital of Zhejiang University School of Medicine and the Second Affiliated Hospital of Zhejiang University School of Medicine. Among the 15 donors, 8 patients were diagnosed with colorectal adenocarcinoma, 1 patient with mucinous adenocarcinoma, 3 patients with adenoma, 2 patients with polyps and 1 with benign tissue at the edge of colorectal tumors. FF tissue samples were snap frozen and stored in liquid nitrogen immediately after surgery and were transported to the proteomics lab within 24h. The colorectal tissue cohort of 15 donors consisted of FFPE and fresh frozen (FF) tissue samples. These samples (1.5×1.5×5 mm^3^ in size) were punched from pathologically confirmed tissue area by Manual Tissue Arrayer MTA-1 (Beecher, US). FF tissue samples were snap frozen and stored in liquid nitrogen immediately after surgery and were transported to the proteomics lab within 24h.

The hepatocellular carcinoma (HCC) cohort and lung adenocarcinoma cohort were collected from Union hospital, Tongji Medical College, Huazhong University of Science and Technology. Sixty-six tissue samples (benign and tumor) from 33 HCC patients were collected within one hour after hepatectomy, then snap frozen and stored at −80 °C. Sixteen tissue samples (matched benign and tumor pairs) from 8 lung adenocarcinoma patients were collected within one hour after pneumonectomy, then snap frozen and stored at −80°C.

The cervical cancer cohort was collected from Tongji Hospital, Tongji Medical College, Huazhong University of Science and Technology. Thirteen FFPE cancerous and benign tissues were obtained from patients with operable cervical cancer.

#### Chinese cancer plasma cohorts

Pooled plasma for building the plasma library was created by mixing plasma (10ul for each patients) from 20 patients from Union Hospital, Tongji Medical College, Huazhong University of Science and Technology. Each of the 20 patients had one of the following hematologic malignancies: acute myelocytic leukemia (AML), acute lymphoblastic leukemia (ALL), chronic myelocytic leukemia (CML), multiple myeloma (MM), myelodysplastic syndrome (MDS) and diffuse large B cell lymphoma (DLBCL). The validation cohort consisted of two groups: 18 clinically healthy volunteers from the Second Affiliated Hospital, Zhejiang University School of Medicine; and 19 patients diagnosed with DLBCL from Union Hospital, Tongji Medical College.

#### Dutch cancer tissue cohorts

The glioblastoma, DLBCL, AML, ALL, cervical, pancreatic and gastric cancer cohorts were collected at Amsterdam UMC/VU medical center, Amsterdam. mirVana aceton precipitations of 19 glioblastoma cancer tissues were pooled by EGFR status (10 wild-type EGFR and 9 mutant (vIII) EGFR samples). Similarly, mirVana aceton precipitations of 27 DLBCL lymphoma patients were pooled by origin (12 samples of neck origin and 17 of non-neck origin). For AML, 2 pools of 2 patient samples each were prepared. For ALL, 14 individual primary ALL cell samples were used, 9 glucocorticoid (GC) resistant and 5 GC sensitive. Cervical cancer tissue lysates of 16 patients were prepared and pooled by subtype (9 SCC and 7 AdCa samples). For pancreatic cancer, individual tissue lysates of 20 patients were used. For gastric cancer, tissues in the form of FFPE material of 10 patients were pooled by tumor percentage (7 with over 50% and 3 with 50% or lower).

The lung cancer cohort was acquired from Amsterdam UMC/VU medical center, Amsterdam and Antoni van Leeuwenhoek hospital/Netherlands Cancer Institute, Amsterdam. Tumor resection samples in the form of FFPE material were collected from 10 lung adenocarcinoma, 10 squamous cell lung carcinoma and 3 large cell lung carcinoma patients and pooled per subtype.

The soft tissue sarcoma cohort was acquired from Antoni van Leeuwenhoek hospital/Netherlands Cancer Institute, Amsterdam. 7 sarcoma and 9 sarcoma metastasis tissues were pooled, respectively.

Prostate and bladder cancer cohorts were acquired from Amsterdam UMC/VU medical center, Amsterdam and Erasmus University Medical Center, Rotterdam. 18 prostate cancer tissues and 9 control tissues in the form of FFPE material were pooled, respectively. In addition, 22 fresh frozen prostate cancer tissues were combined to 2 pools of 11 samples each. 10 bladder cancer tissues in the form of FFPE material were pooled in 2 pools of 5 samples each.

The CRC and triple-negative breast cancer (TNBC) cohorts were collected at Erasmus University Medical Center, Rotterdam. For CRC, 2 pools were made per CMS subtype (CMS1, 2, 3 and 4), whereby each pool contained tissue lysates of 5 patients. For TNBC, 2 pools of 23 and 24 patient tissues each were used.

#### Singapore thyroid cancer cohort

The thyroid tissue cohort was kindly provided by National Cancer Centre, Singapore. 105 FFPE thyroid tissue punches from 63 patients were included in this study. The cohort is composed of 5 patients with normal thyroid, 28 with multinodular goiter, 10 with follicular thyroid adenoma, 5 with follicular thyroid carcinoma and 15 with papillary thyroid carcinoma.

### Pre-treatment and de-crosslinking of FFPE tissue samples

About 1 mg of FFPE tissue was first dewaxed three times by heptane, then rehydrated in a gradient of 100%, 90%, 75% ethanol. The partly rehydrated samples were then transferred into microtubes (PBI, MA, USA) and soaked in 0.1% formic acid (FA) for complete rehydration and acidic hydrolysis for 30 min, under shaking at 600 rpm, 30°C. The thus treated FFPE samples were washed using 0.1 M Tris-HCl (pH 10.0) by gentle shaking and spinning down. The supernatant was discarded. 15 μL of 0.1 M Tris-HCl (pH 10.0) was added to cover tissues and the suspension was boiled at 95 °C for 30 min for basic hydrolysis under gentle shaking. Subsequently the sample was fast cooled to 4°C, topped with 25 μL of lysis buffer containing 6M urea and 2M thiourea, 0.1mM NH_4_HCO_3_ (pH 8.5), and subjected to PCT-assisted tissue lysis and digestion.

### PCT-assisted tissue lysis and digestion

About 1mg of de-crosslinked FFPE tissue or pre-washed FF tissue was mixed with 35μL lysis buffer containing 6M urea and 2M thiourea, 0.1mM NH_4_HCO_3_ (pH 8.5) in microtubes and capped with micropestles (PBI, MA, USA). Alternatively, if the proteins were extracted for later 1D SDS-page separation, 1% SDS in Milli-Q water was used instead of urea/thiourea lysis buffer. Tissues were lysed in a barocycler NEP2320-45k (Pressure BioSciences Inc.) at the PCT scheme of 30s high pressure at 45kpsi plus 10s ambient pressure, oscillating for 90 cycles at 30°C. Extracted proteins were reduced and alkylated by incubating with 10mM Tris(2-carboxyethyl) phosphine (TCEP) and 20mM iodoacetamide (IAA) at 25 °C for 30 min, in darkness, by gently vortexing at 800 rpm in a thermomixer. Afterwards, proteins were digested by Lys-C (Hualishi Beijing; enzyme-to-substrate ratio, 1:40) using the PCT scheme of 50 s high pressure at 20 kpsi plus 10 s ambient pressure, oscillating for 45 cycles at 30°C. This was followed by a tryptic digestion step followed (Hualishi Beijing; enzyme-to-substrate ratio, 1:50) using the PCT scheme of 50 s high pressure at 20kpsi plus 10s ambient pressure, oscillating for 90 cycles at 30°C. Finally, 15 μL of 10% trifluoroacetic acid (TFA) was added to each tryptic digest to quench the enzymatic reaction (final concentration of 1% TFA). Peptides were purified by BioPureSPN Midi C18 columns (The Nest Group Inc., Southborough, MA) according to the manufacturer’s protocol. Peptide eluates were then dried under vacuum (LABCONCO CentriVap, Kansas, MO). Dry peptides were dissolved in 20 μL of water containing 0.1% FA and 2% ACN (acetonitrile) (all MS grade). Peptide concentration was measured using ScanDrop^2^ (AnalytikJena, Beijing, China) at A280.

### 1D SDS-PAGE separation at protein level for building DDA library

SDS-PAGE separation and peptide preparation in Jimenez lab, the Netherlands: Tissues were lysed in 1× reducing NuPAGE LDS sample buffer (Invitrogen, Carlsbad, CA), sonicated in a Branson cup-type digital sonifier, centrifuged, and heated for 5 minutes at 95°C. Protein lysates were separated on precast 4-12% gradient gels using the NuPAGE SDS-PAGE system (Invitrogen, Carlsbad, CA). Following electrophoresis, gels were fixed in 50% ethanol/3% phosphoric acid solution and stained with Coomassie R-250. Subsequently, gel lanes were cut into 10 bands and each band was cut into ~1 mm^3^ cubes. The gel cubes from each band were transferred into a well of a 96-well filter plate (Eppendorf, Hamburg, Germany) and were washed in 50 mM NH_4_HCO_3_ and 2x 50 mM NH_4_HCO_3_/50% ACN. Subsequently, gel cubes were reduced for 60 min in 10 mM dithiothreitol (DTT) at 56°C and alkylated for 45 min in 50 mM IAA (both Sigma, St Louis, MO) in the dark, at room temperature. After washing in 50 mM NH_4_HCO_3_ and 2x 50 mM NH_4_HCO_3_ /50% ACN, the gel cubes were dried for 10 min in a vacuum centrifuge at 60°C and subsequently incubated in 50 μl 6.25 ng/μL sequence-grade trypsin (Promega, Madison, WI) in 50 mM NH_4_HCO_3_ at room temperature overnight. Peptides from each gel band were extracted once using 150 μL 1% FA, and twice using 150 μL 5% FA/50%ACN and were pooled in a 96-deep-well plate and centrifuged to dryness at 60°C in a vacuum centrifuge and stored at −20°C. Dried peptide extracts were dissolved in 25μL loading solvent (0.5% TFA in 4% ACN) containing 2.5 injection equivalent (IE) iRT retention time peptide standard (Biognosys, Schlieren, CH). 5 μL of peptide extract containing 0.5 IE iRT peptides was injected into the nanoLC system.

SDS-PAGE separation and peptide preparation in Guo lab, China: About 200-300 μg of protein was mixed with 3× SDS sample loading buffer (GenScript Biotech, China) supplemented with 150 mM DTT, and the mixture was boiled at 95°C for 5 min.1D gel electrophoresis was performed using 4-12% gradient SDS-PAGE after which the gel was removed, washed first with distilled water and then with the fixing buffer (50% (v/v) ethanol in water with 5% (v/v) acetic acid) at room temperature for 15 min with gentle agitation to remove excessive SDS. The fixed and washed gel was stained in Coomassie blue for around 1 h with gentle agitation, and then de-stained until the background was clear and protein bands were visible. The gel was rehydrated in distilled water at room temperature for 30 min with gentle agitation. Ten protein bands to cover each lane were cut out and further cut into ca 1 × 1 mm^2^ pieces, followed by reduction with 10 mM TCEP in 25mM NH_4_HCO_3_ at 25°C for 1 h, alkylation with 55 mM IAA in 25 mM NH_4_HCO_3_ solution at 25°C in the dark for 30 min, and sequential digestion with trypsin at a concentration of 12.5 ng/mL at 37°C overnight (1^st^ digestion for 4hrs and 2^nd^ digestion for 12hrs). Tryptic-digested peptides from gel pieces were extracted three times using 50% ACN/5% FA and dried under vacuum. Dry peptides were purified by Pierce C18 Spin Tips (Thermo Fisher, USA).

### Preparation and fractionation of plasma protein samples

Venous blood of each patient was collected in EDTA and anticoagulation proceeded for 9 hours. Plasma samples obtained by centrifugation were transferred to a new set of 1.5 mL Eppendorf tubes and stored at 4°C. Samples were cold-transported from the hospital to the proteomics lab within 36 h at 4°C. Samples were centrifuged again at 300g for 5min at 4°C to remove cells and the supernatants were further centrifuged at 2500g for 15min at 4°C to remove cell debris and platelets. The final supernatants were stored at −80°C for further protein extraction and in solution digestion.

To remove very high abundant plasma proteins in this study, whole plasma peptides were further extensively fractionated by several methods such as SDS-PAGE separation, antibody-depletion of high abundant proteins and exosome isolation.

For SDS-PAGE fractionation, the entire gel was cut into 12 thin gel rows, of which four rows with heavily stained protein bands (3 adjacent bands between 45-75 kD, and a band between 25 and 35 kD) were picked out for depletion of high abundant proteins. Each of the other 8 rows was subjected to in-gel digestion as described above. We also used High Select Top 14 Abundant Protein Depletion Resin spin columns (Thermo Scientific, A36370) to deplete high abundance proteins in plasma samples according to the manufacturer’s instructions; and further fractionated and digested the depleted plasma proteins by 1D SDS-PAGE.

To obtain the enriched exosome fraction, an aliquot of 200 μL plasma was taken after centrifuging venous blood for 10 min at 3000 g, 4°C. The exosome pellet was collected after ultracentrifugation at 160,000g, 4°C for 12h and resuspended in cold phosphate-buffered saline for washing. Resuspended exosomes were further centrifuged at 100,000g, 4°C for 70 min. The pellet was collected and redissolved in 150 μL of 2% SDS. The exosome fraction in 2% SDS was subjected to PCT-assisted sample lysis, undergoing 60 cycles at 20°C, with 45 k p.s.i. for 50s and atmosphere pressure for 10 s. After lysis, the exosome protein mixture was precipitated with 80% cold acetone at −20°C for 3h and the suspension was centrifuged at 12,500 g, 4°C for 15 min to collect the protein pellet. The protein pellet was redissolved with 200 μL of 1% SDS, followed by SDS-PAGE separation and subsequent in-gel digestion. Each exosome protein sample was cut into three fractions and digested as described above.

### Strong cation-exchange (SCX) fractionation at peptide level for building DDA library

The SCX solid phase extraction (SPE) cartridge (Thermo Scientific, # 60108-421) was conditioned first according to the manufacturer’s protocol. For SCX fractionation, about 1mg peptides were dissolved in 1 mL of 5 mM KH_2_PO_4_/25%ACN (pH = 3.0), then the peptide solution was loaded onto the well-conditioned SCX SPE cartridge. The cartridge was then rinsed with 5mM KH_2_PO_4_/25%ACN (pH = 3.0). Finally, six peptide fractions were collected by eluting the cartridge with 1.5 mL increments of increasing KCl concentration in 5mM KH_2_PO_4_/25%ACN, i.e. 50 mM, 100 mM, 150 mM, 250 mM, 350 mM, and 500 mM. Each fraction was collected and vacuumed to dryness. Dry peptides and precipitated salts were redissolved in 200μL of 0.1% TFA and subjected to further C18 desalting by BioPureSPN Midi SPE (Nest Group, Cat # HEM S18V).

### DDA data acquisition in Jimenez lab

547 DDA raw data files were generated at Jimenez lab. All peptides were prepared via SDS-PAGE fractionation and in-gel digestion. Peptides were separated by an Ultimate 3000 nanoLC-MS/MS system (Dionex LC-Packings, Amsterdam, The Netherlands) equipped with a 40 cm × 75 μm ID fused silica column custom packed with 1.9 μm 120Å ReproSil Pur C18 aqua (Dr Maisch GMBH, Ammerbuch-Entringen, Germany). After injection, peptides were trapped at 10μL/min on a 10mm × 100 μm ID trap column packed with 5 μm 120Å ReproSil Pur C18 aqua in 0.1% formic acid. Peptides were separated at 300 nL/min in a 10–40% linear gradient (buffer A: 0.1% formic acid (Fischer Scientific), buffer B: 80% ACN, 0.1% formic acid) in 90 min (130 min inject-to-inject). Eluting peptides were ionized at a potential of +2 kV into a Q Exactive mass spectrometer (Thermo Fisher, Bremen, Germany). Intact masses were measured at resolution 70,000 (at m/z 200) in the orbitrap using an AGC target value of 3E6 charges and an S-lens setting of 60. The top 10 peptide signals (charge-states 2+ and higher) were submitted to MS/MS in the HCD (higher-energy collision) cell (1.6 amu isolation width, 25% normalized collision energy). MS/MS spectra were acquired at resolution 17,500 (at m/z 200) in the orbitrap using an AGC target value of 1E6 charges, a max injection time (IT) of 80ms and an underfill ratio of 0.1%. Dynamic exclusion was applied with a repeat count of 1 and an exclusion time of 30 s.

### DDA Data acquisition in Guo Lab

549 DDA raw data files were generated at Guo lab. Biognosys-11 iRT peptides (Biognosys, Schlieren, CH) were spiked into peptide samples at the final concentration of 10% prior to MS injection for RT calibration. Peptides were separated by Ultimate 3000 nanoLC-MS/MS system (Dionex LC-Packings, USA) equipped with a 15 cm × 75μm ID fused silica column packed with 1.9μm 100Å C18. After injection, peptides were trapped at 6 μL/min on a 20 mm × 75 μm ID trap column packed with 3 μm 100 Å C18 aqua in 0.1% formic acid. Peptides were separated along a 120min 3–25% linear LC gradient (buffer A: 2% ACN, 0.1% formic acid (Fisher Scientific), buffer B: 98% ACN, 0.1% formic acid) at the flowrate of 300 nL/min (148 min inject-to-inject). Eluting peptides were ionized at a potential of +1.8 kV into a Q-Exactive HF mass spectrometer (Thermo Fisher, Bremen, Germany). Intact masses were measured at resolution 60,000 (at m/z 200) in the orbitrap using an AGC target value of 3E6 charges and a S-lens setting of 50. The top 20 peptide signals (charge-states 2+ and higher) were submitted to MS/MS in the HCD (higher-energy collision) cell (1.6 amu isolation width, 27% normalized collision energy). MS/MS spectra were acquired at resolution 30,000 (at m/z 200) in the orbitrap using an AGC target value of 1E5 charges, a max IT of 80ms and an underfill ratio of 0.1%. Dynamic exclusion was applied with a repeat count of 1 and an exclusion time of 30 s.

### DIA data acquisition in Guo lab

The LC configuration for DIA data acquisition is as the same as for DDA data acquisition with slight modifications. Biognosys-11 iRT peptides (Biognosys, Schlieren, CH) were spiked into peptide samples at the final concentration of 10% prior to MS injection for RT calibration. Peptides were separated at 300 nL/min in a 3–25% linear gradient (buffer A: 2% CAN, 0.1% formic acid (Fischer Scientific), buffer B: 98% ACN, 0.1% formic acid) in 45 min (68 min inject-to-inject). Eluting peptides were ionized at a potential of +1.8 kV into a Q-Exactive HF mass spectrometer (Thermo Fisher, Bremen, Germany). A full MS scan was acquired analyzing 390-1010 m/z at resolution 60,000 (at m/z 200) in the orbitrap using an AGC target value of 3E6 charges and maximum IT 80ms. After the MS scan, 24 MS/MS scans were acquired, each with a 30,000 resolution at m/z 200, AGC target 1E6 charges, normalized collision energy was 27%, with the default charge state set to 2, maximum IT set to auto. The cycle of 24 MS/MS scans (center of isolation window) with three kinds of wide isolation window are as follows (m/z): 410, 430, 450, 470, 490, 510, 530, 550, 570, 590, 610, 630, 650, 670, 690, 710, 730, 750, 770, 790, 820, 860, 910, 970. The entire cycle of MS and MS/MS scans acquisition took roughly 3s and was repeated throughout the LC/MS/MS analysis.

### DIA Data analysis using OpenSWATH and TRIC

Briefly, DIA raw data files were converted in profile mode to mzXML using msconvert and analyzed using OpenSWATH (2.0.0) [14] as described [13]. Retention time extraction window was 600 seconds, and m/z extraction was performed with 0.03Da tolerance. Retention time was then calibrated using both SiRT and CiRT peptides. Peptide precursors that were identified by OpenSWATH and pyprophet with d_score >0.01 were used as inputs for TRIC [51]. For each protein, the median MS2 intensity value of peptide precursor fragments which were detected to belong to the protein was used to represent the protein abundance.

### Terms for protein identifications

In this paper, the term “protein group” indicates a group of proteins sharing identified peptides appeared in all the protein members. Proteins identified from SwissProt protein sequence database (i.e. one manually inspected protein sequence per gene symbol, excluding isoforms, splicing variants and theoretical protein sequences) are called “SwissProt proteins”. The proteotypic protein refers to a protein which is identified by proteotypic peptides which only appear in one SwissProt protein sequence.

### Validation of representative proteins using parallel reaction monitoring (PRM)

PRM quantification strategy was used to further validate proteins that were measured by DIA quantification above. Biognosys-11 iRT peptides (Biognosys, Schlieren, CH) were spiked into peptide samples at the final concentration of 10% prior to MS injection for RT calibration. Peptides were separated at 300 nL/min along a 60min 7–35% linear LC gradient (buffer A: 20% ACN, 0.1% formic acid; buffer B: 20% ACN, 0.1% formic acid). The Orbitrap Fusion Lumos Tribrid mass spectrometer was operated in the MS/MS mode with time-scheduled acquisition for 100 peptides in a +/− 5 min retention time window. The individual isolation window was set at 1.2 Th. The full MS mode was measured at resolution 60,000 at m/z 200 in the Orbitrap, with AGC target value of 4E5 and maximum IT of 50ms. Target ions were submitted to MS/MS in the HCD cell (1.2 amu isolation width, 30% normalized collision energy). MS/MS spectra were acquired at resolution 30,000 (at m/z 200) in the Orbitrap using AGC target value of 1E5, a max IT of 100ms.

## AVAILABILITY

Computational pipeline as a Docker container and DPHL as .tsv flat file initiative is available in the OneDrive website (https://westlakeu-my.sharepoint.com/:f:/g/personal/zhutiansheng_westlake_edu_cn/En-CNWLzaAxCja-L8Jze-6cBLHi7FTeIJNLnNcRMQacH5g?e=WOKizE)

## ACCESSION NUMBERS

All the DDA files, DIA-MS Data files, original peptides, and protein results are deposited in iProX; the Project ID is IPX0001400000 and can be accessed via http://www.iprox.org/page/PSV023.html;?url=1542762994917ZL13. All data and codes will be publicly released upon publication.

## SUPPLEMENTARY DATA

Supplementary Data are available at NAR online.

## ACKNOWLEDGEMENTS

The authors thank all collaborators who participated in the procurement of the clinical specimens.

