## Supplementary Note for "DPHL: A pan-human protein mass spectrometry library for robust biomarker discovery"

**Supplementary Figure 1**

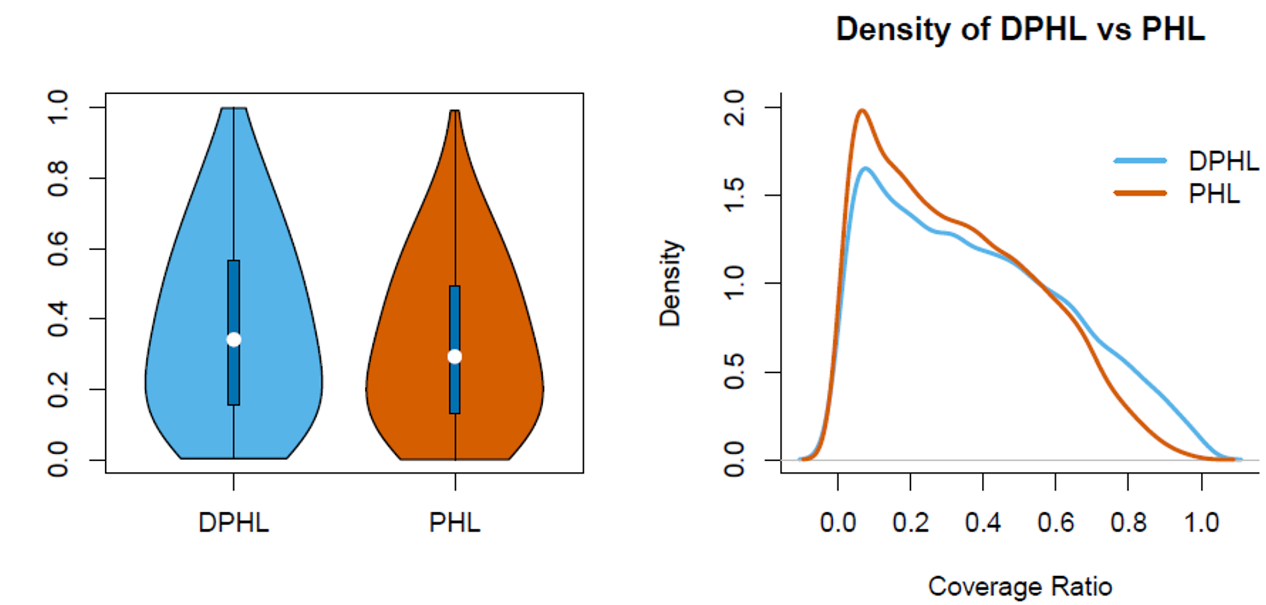

Density

Sequence Coverage Ratio

Sequence Coverage Ratio

**Supplementary Figure S1. Comparison of protein sequence coverage between DPHL and PHL.** Violin plot (left panel) and density plot (right panel) of protein sequence coverage for the proteins identified in the two libraries.

**Supplementary Figure S2A**

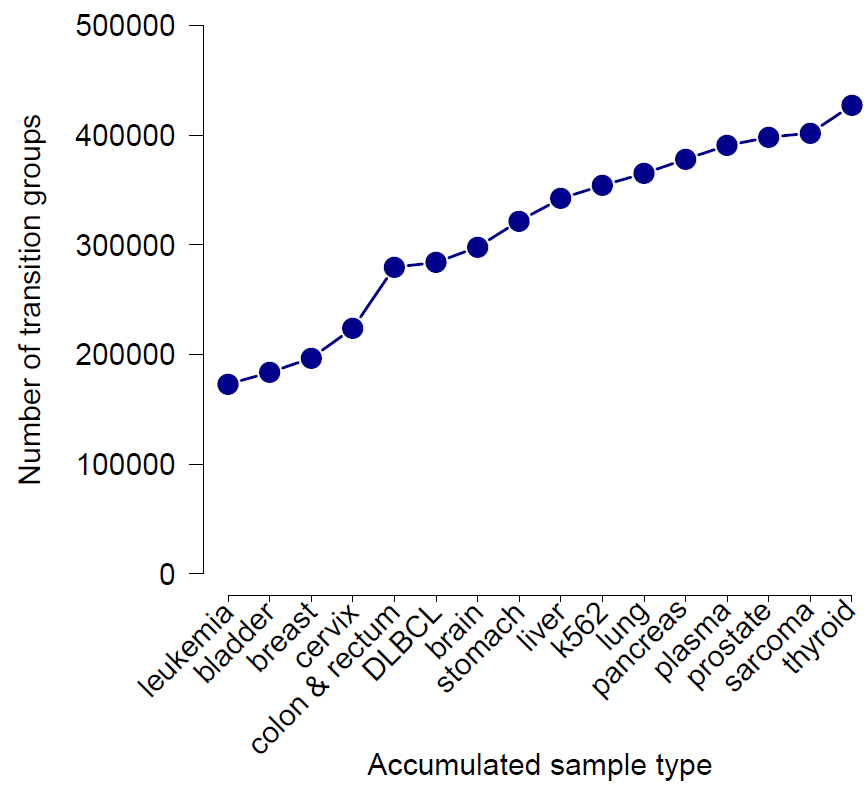

**Supplementary Figure S2A. Number of transition groups in accumulated data sets.** The number of transition groups in DPHL increases cumulatively after adding data sets from various tissue types.

**Supplementary Figure S2B**

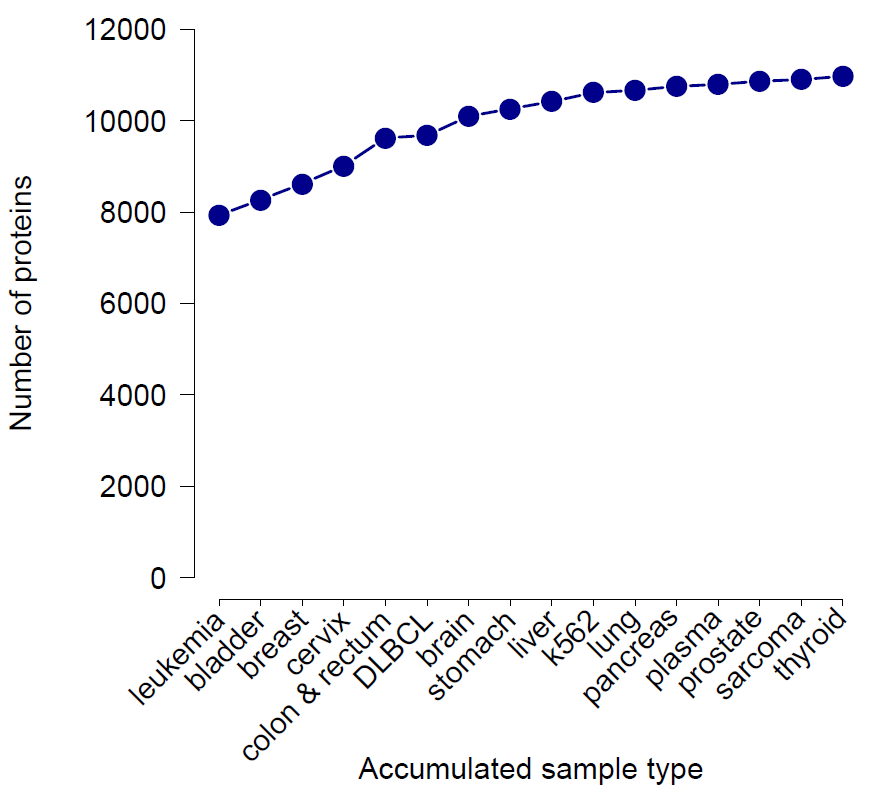

**Supplementary Figure S2B. Cumulative count of proteins in the DPHL library.** The number of proteins saturates as the number of raw data files increase.

**Supplementary Figure S2C**

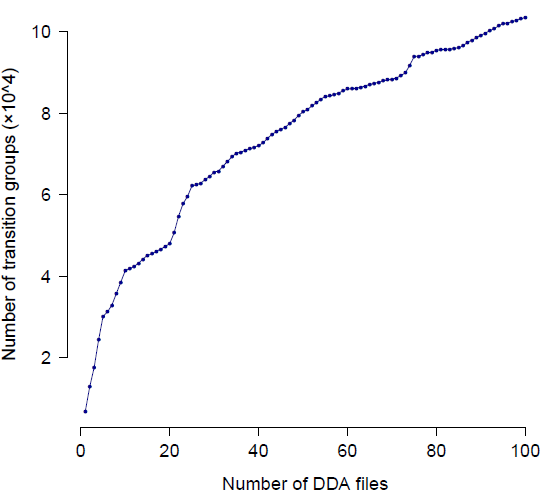

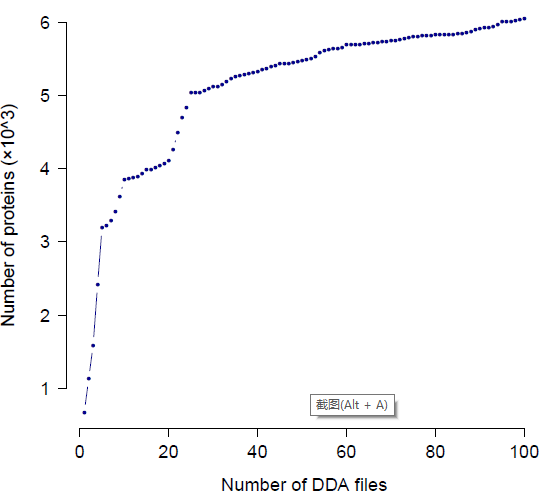

**Supplementary Figure S2C. Cumulative count of peptides and proteins identified from the pancreas tissue samples based on 100 DDA files.** Upper panel: number of transition groups (*i.e.* peptide precursors); lower panel: number pf proteins.

**Supplementary Figure S2D**

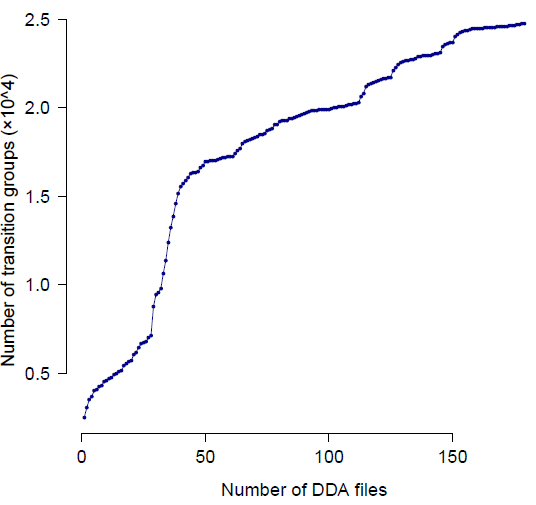

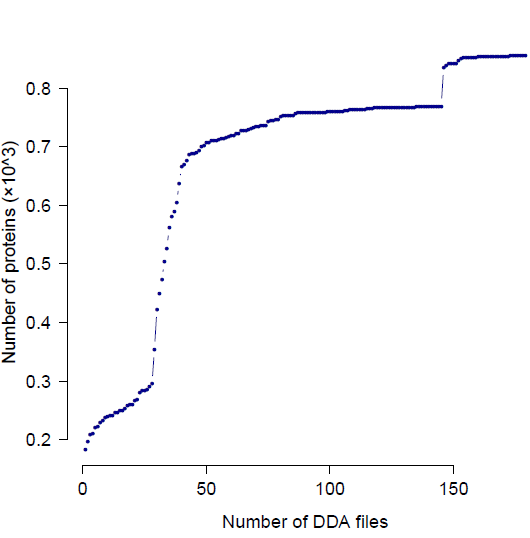

**Supplementary Figure S2D. Cumulative count of peptides and proteins identified from the 179 plasma samples.** Upper panel: number of transition groups (i.e. peptide precursors); lower panel: number pf proteins.

**Supplementary Figure S3**

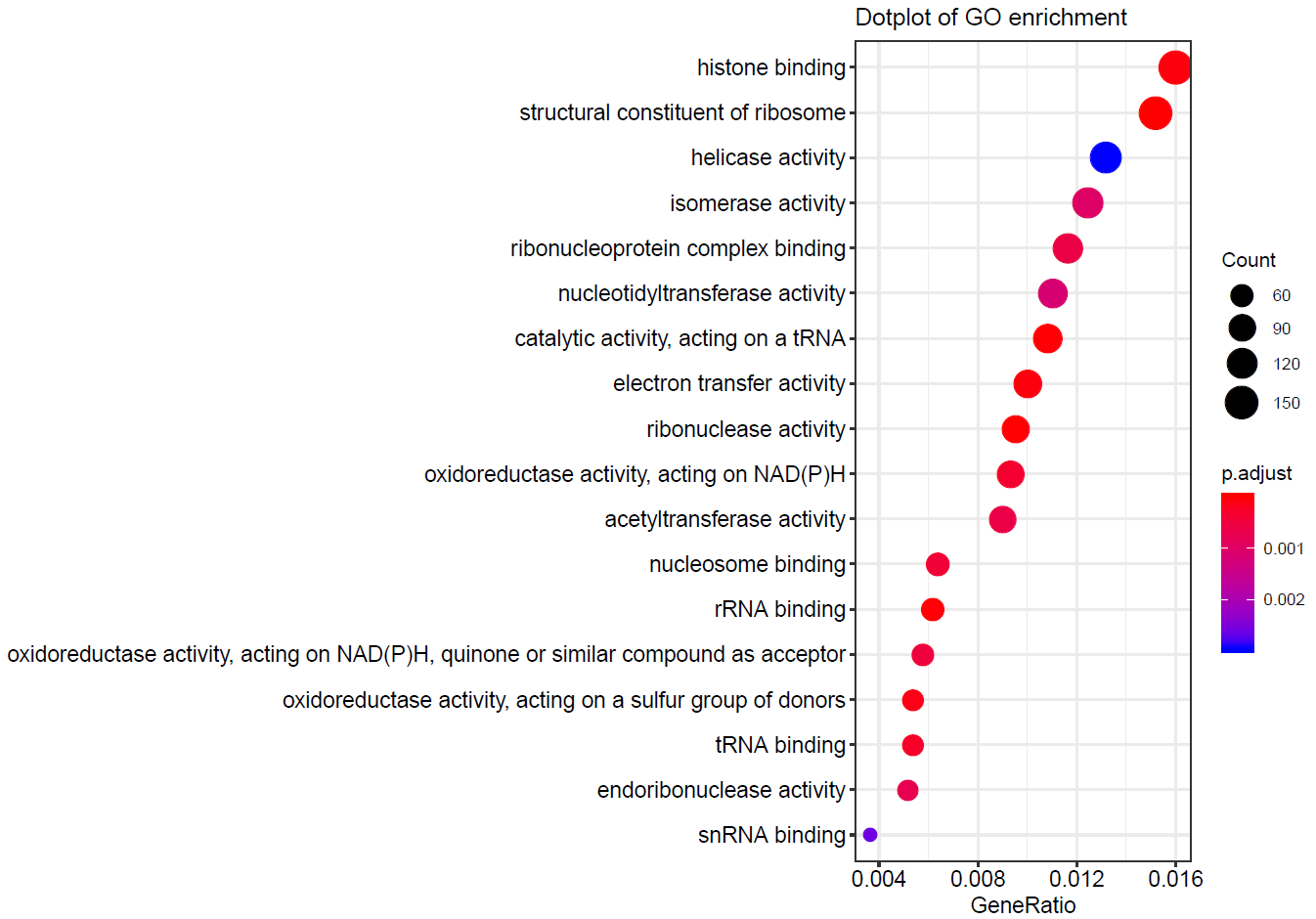

A

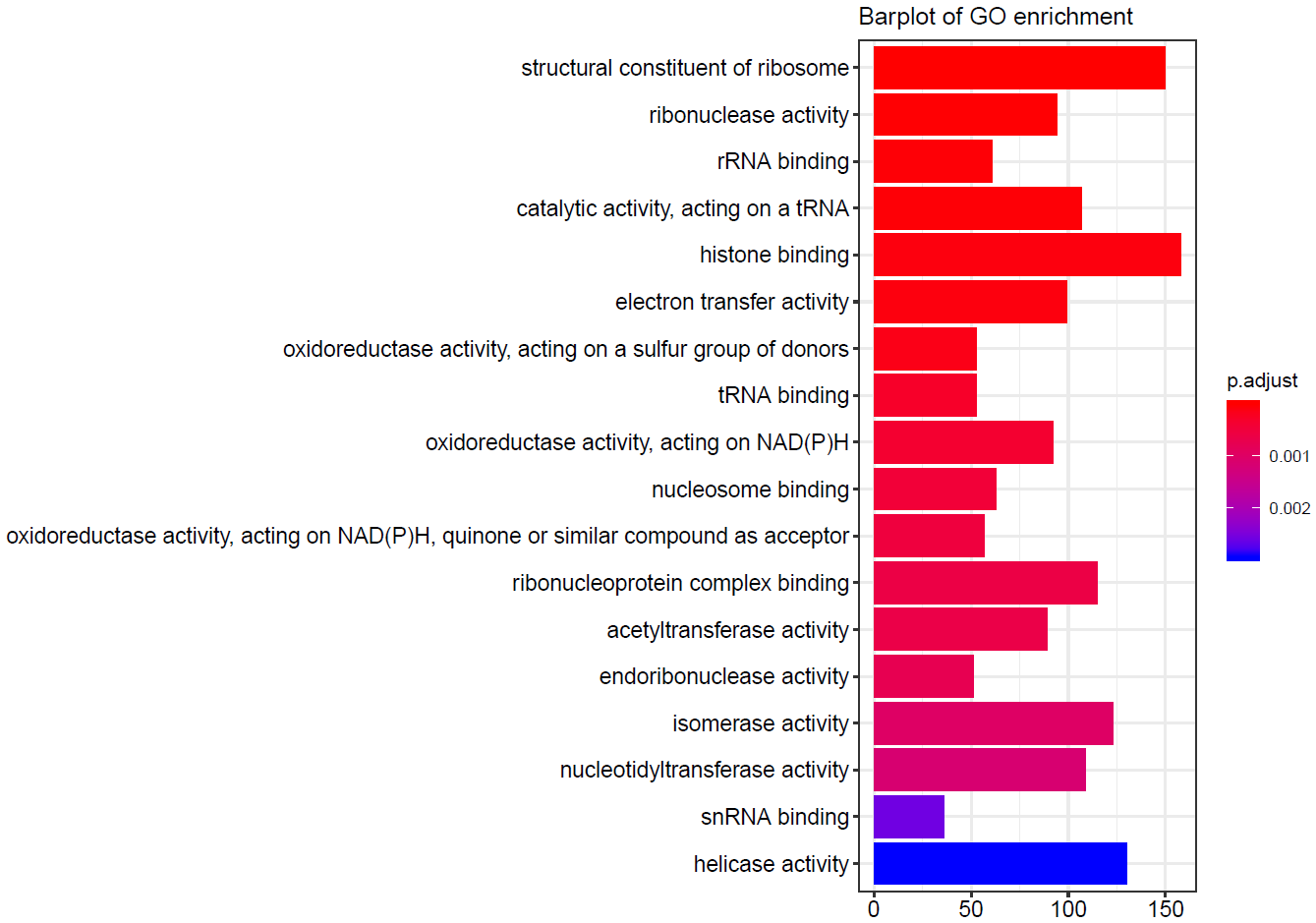

B

**Supplementary Figure S3. GO annotation of all proteins identified in DPHL.** 18 gene ontology groups are shown after GO molecular function enrichment analysis. (**A**) Size of circle: count of proteins; color of circle: adjust *P* values. The X-axis label “GeneRatio” means the ratio of number of genes in DPHL to all the genes with the respective GO term. (**B**) Length of bar: number of proteins; color of bar: adjust *P* values.

**Supplementary Figure S4**

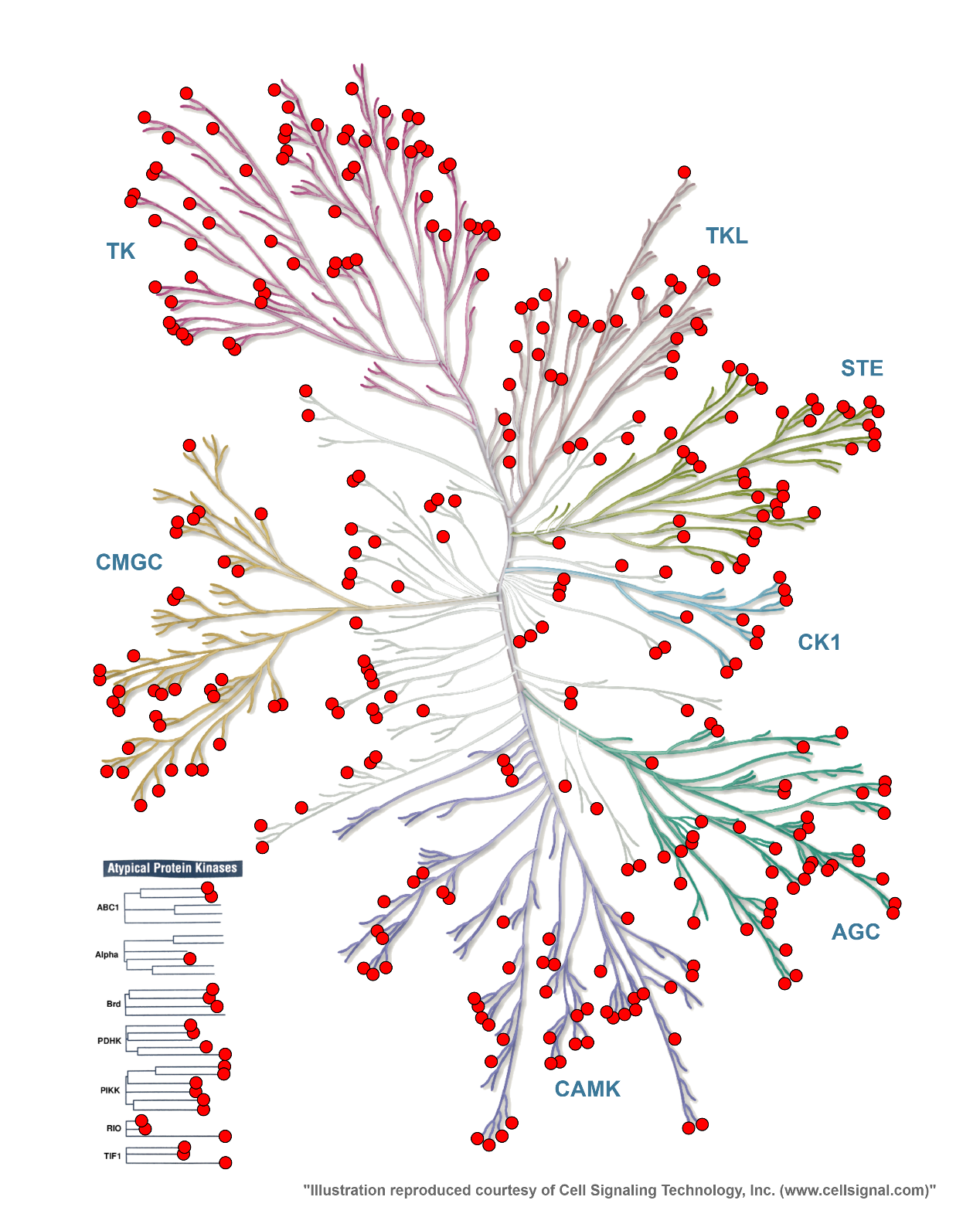

**Supplementary Figure S4. KinMap of proteins in the DPHL library.** The KinMap contains eight typical groups (AGC, CAMK, CK1, CMGC, STE, TK, TKL, Other). The DPHL covers almost the entire branches of kinome tree. 347 proteins (red dots) in the DPHL library are detected in the KinMap tree. The lower left corner shows the proteins from DPHL in the atypical kinase families.

**Supplementary Figure S5**

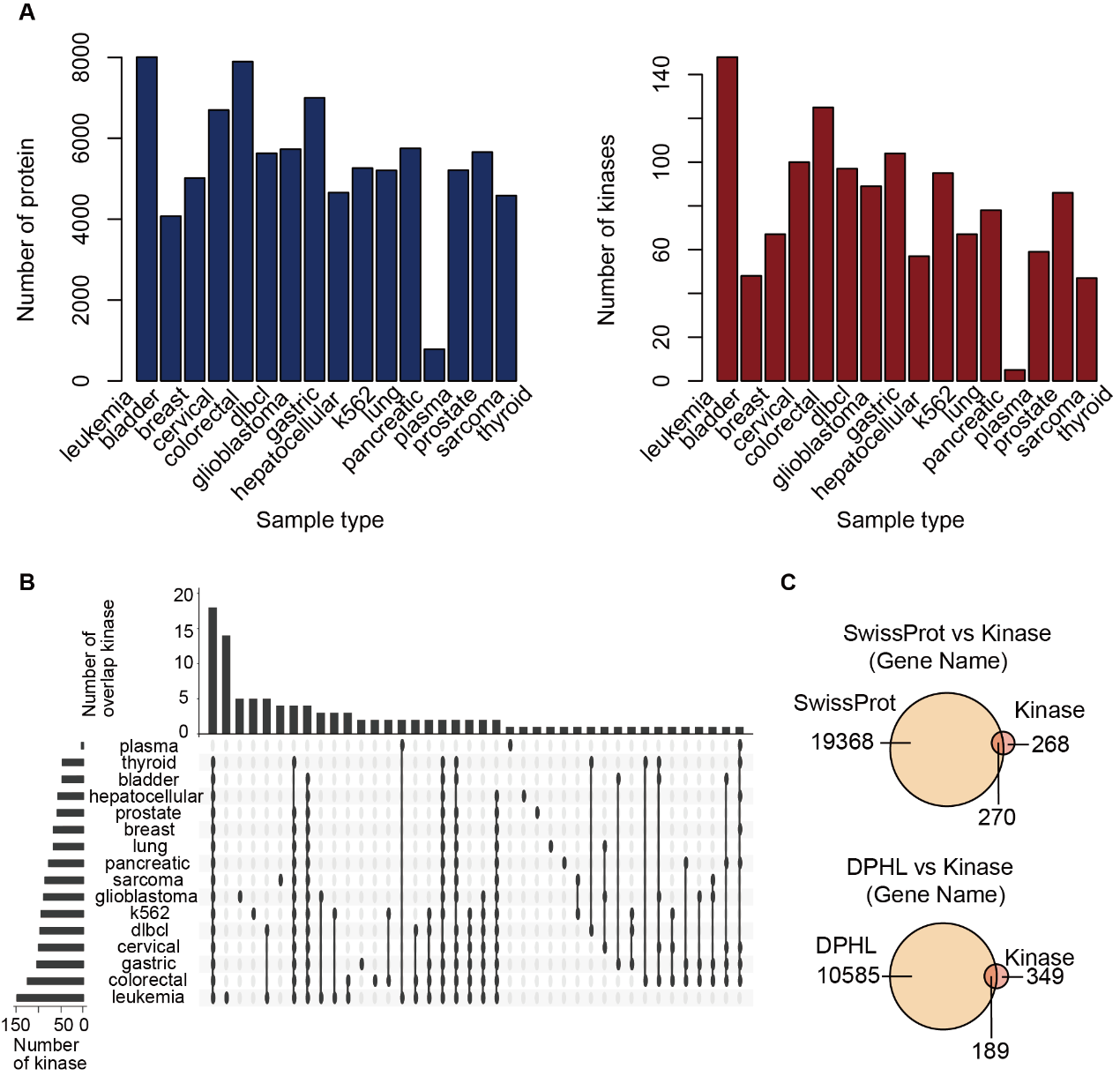

**Supplementary Figure S5. Protein kinases identified in the DPHL library. (A)** The bar charts show the number of proteins identified (left blue) and the kinases (right red) in the DPHL library. **(B)** shows the overlap of kinases for different tissue types. **(C)** The Venn diagrams show the comparison of DPHL with SwissProt and the kinome. DPHL covers most kinase in SwissProt ((198 / 270) × 100%= 70%).

**Supplementary Figure S6**

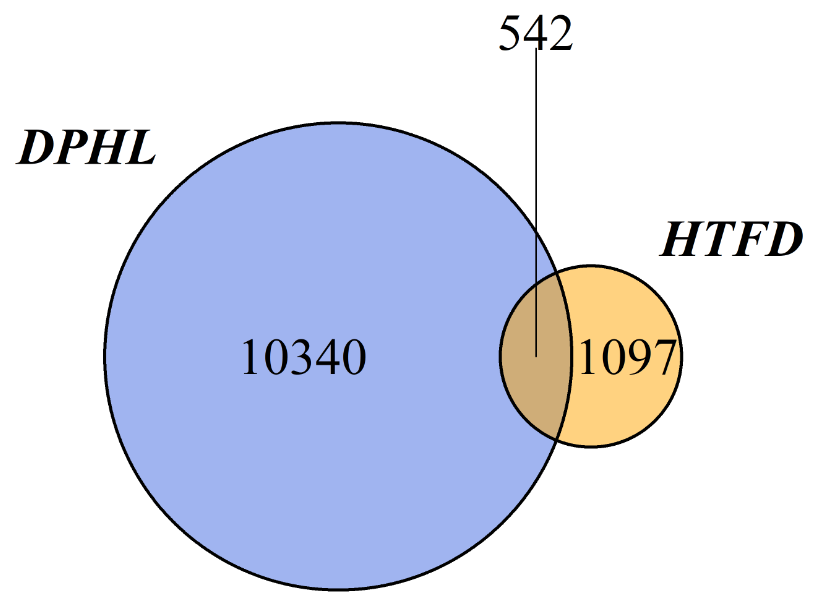

**Supplementary Figure S6.** 542 TFs proteins were identified in DPHL, using the Human Transcription Factors database (HTFD) as a reference.

**Supplementary Figure S7**

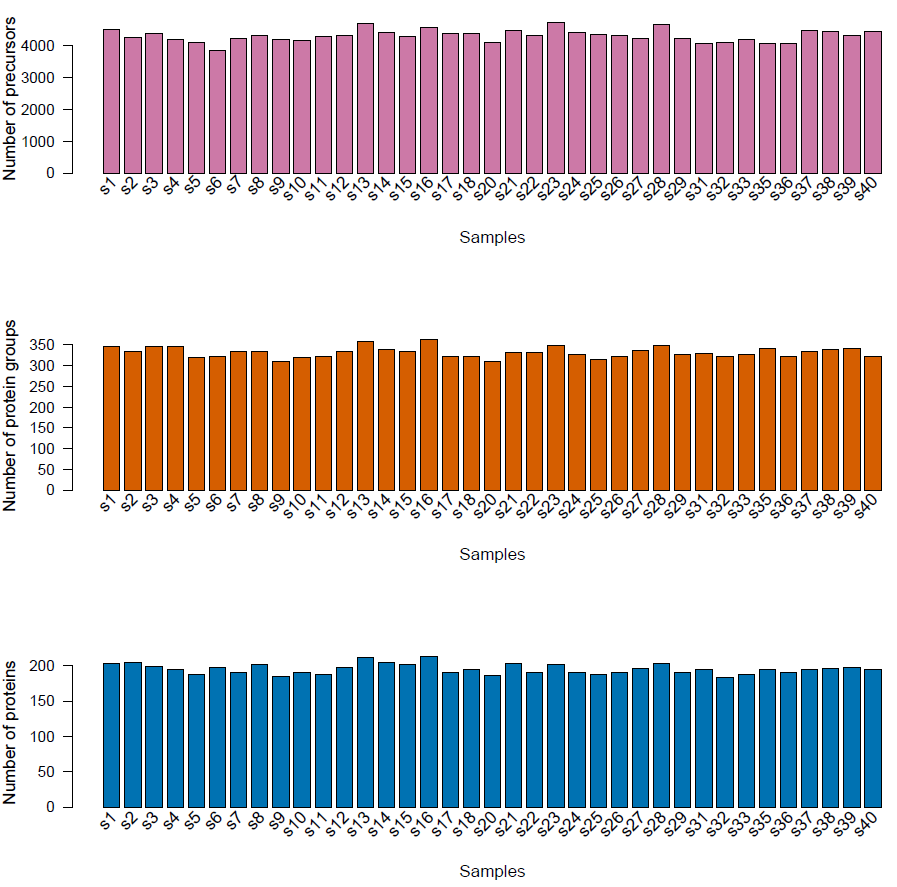

**Supplementary Figure S7.** Number of peptide precursors (mean: 4,313, min: 3,855; max: 4,728), protein groups (mean: 332, min: 309; max: 363) and proteotypic proteins (mean: 195, min: 184; max: 213) identified in the 37 DLBCL plasma samples.

**Supplementary Figure S8**

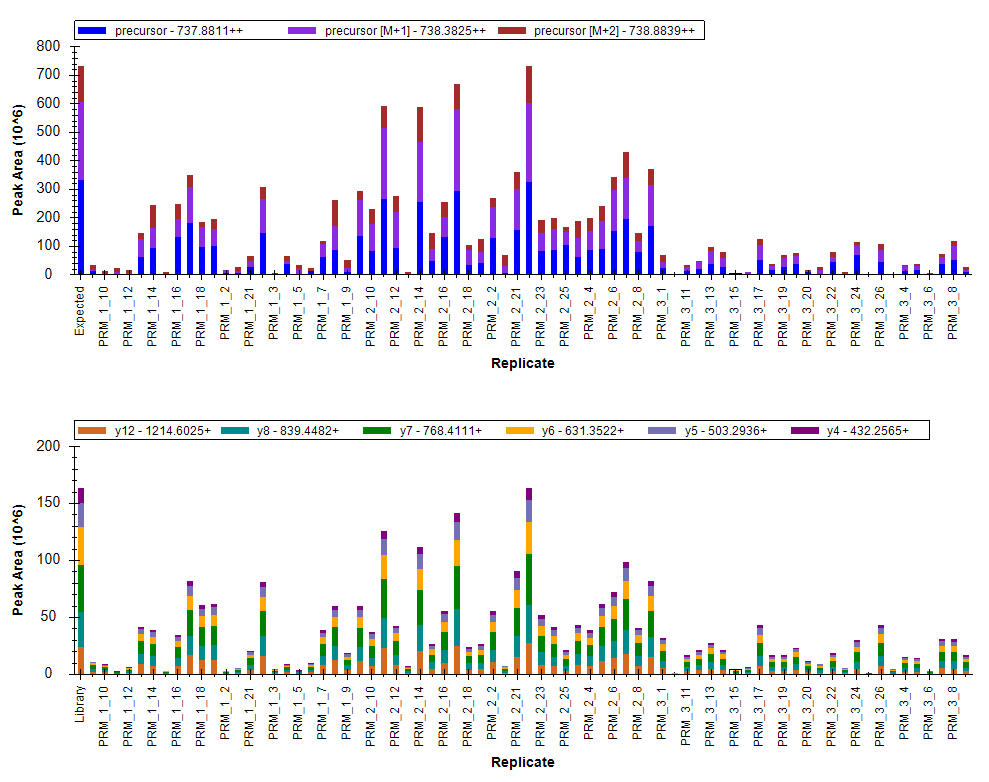

**Supplementary Figure S8. Skyline quantified the relative abundance of TPP1 (O14773) from the peak area information of its best flying peptides.** One representative example: LFGGNFAHQASVAR (m/z 737.88) of TPP1 (O14773) across 73 prostate samples. The upper panel shows the precursor quantification at MS1 level and the lower panel depicts the fragment information at MS2 level.

**Supplementary Figure S9**

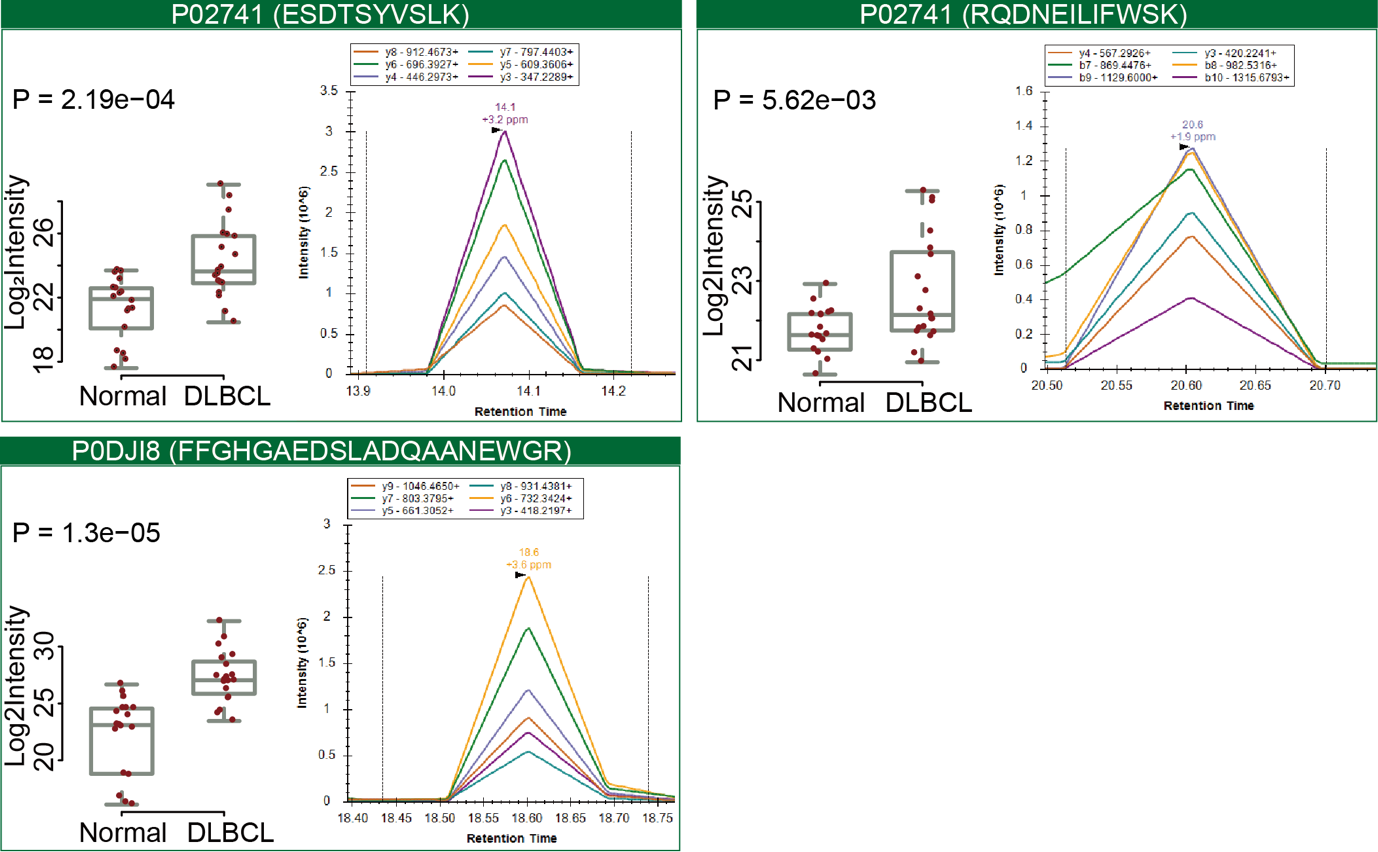

**Supplementary Figure S9**. PRM validation of three peptides in 37 plasma samples. In each box, the left panel shows the log_2_ intensity of the respective peptides across 37 plasma samples, while the right panel depicts a representative peak group for the peptide from s8 sample. P values are computed using Student’s *t* test.

**Supplementary Figure S10**

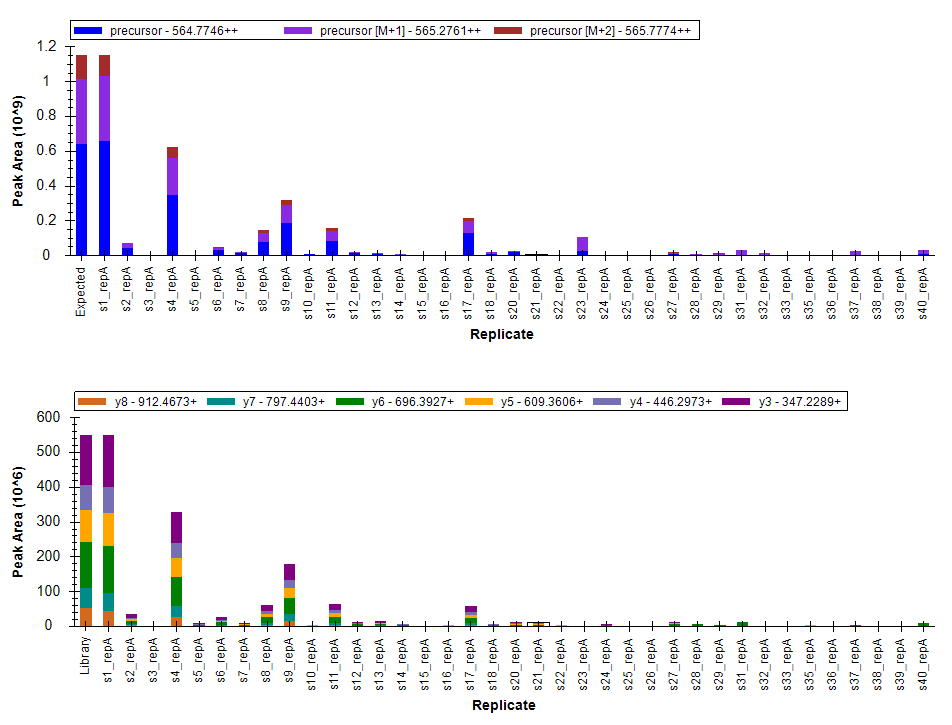

**Supplementary Figure S10. The peak area for peptide ESDTSYVSLK (m/z 564.77) of CRP (P02741) across 37 plasma samples.** The upper panel shows the precursor quantification and the lower panel depicts the fragment information.

**Supplementary Note 2. Comparison of DDA files acquired from the Guo lab and the Jimenez lab.**

We compared the consistency of DDA files acquired from the Guo lab and the Jimenez lab. Although the two laboratories both employed QE-HF instrument for the project, however there are potentially difference in data due to biological heterogeneity and instrument difference. We selected two sub data sets from stomach since both laboratories acquired DDA data for these two types of tissues. The Guo lab acquired 70 DDA files from the Chinese stomach cancer tissue samples, while the Jimenez lab obtained 20 DDA files from the Dutch patients. The intention here was not to shoot the same samples but to mimic the real world situation and compare the biological data from two laboratories.

The Guo lab identified more peptide precursors, peptides and proteins than the Jimenez lab , however, the two sub data sets shared 26,822 peptide precursors, 24,461 peptides and 4,739 proteins (Figure N2A). We also found that most of the top 6 fragments are shared by the two sub data sets (Figure N2B).

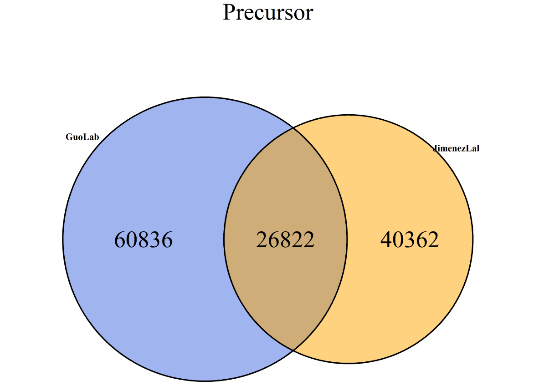

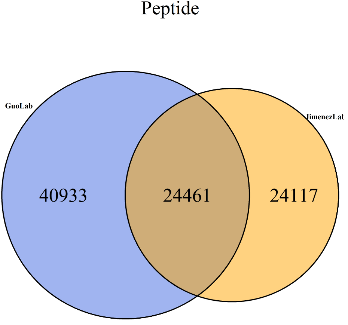

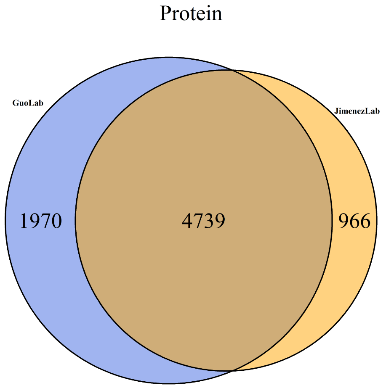

**Figure N2A. Venn diagrams for number of peptide precursors, peptides and proteins identified in the Guo lab and the Jimenez lab**.

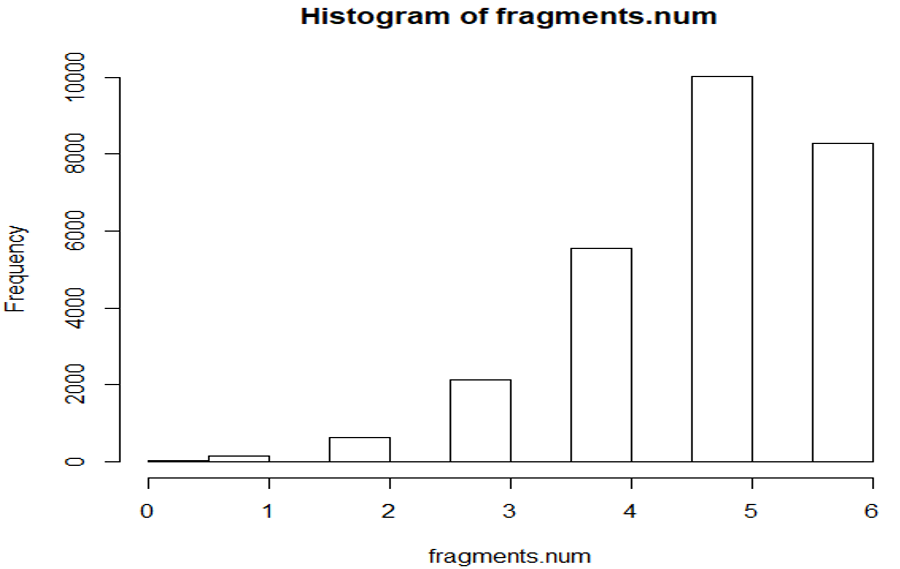

**Figure N2B. Shared top 6 fragments identified in the Guo lab and the Jimenez lab.**

We then compared the iRT-calibrated retention time, the Pearson correlation coefficient reach 0.99 (Figure N2C). The correlation got worse when the iRT values are higher than 100, due to the slightly shorter effective gradient in the Guo lab to increase sample throughput.

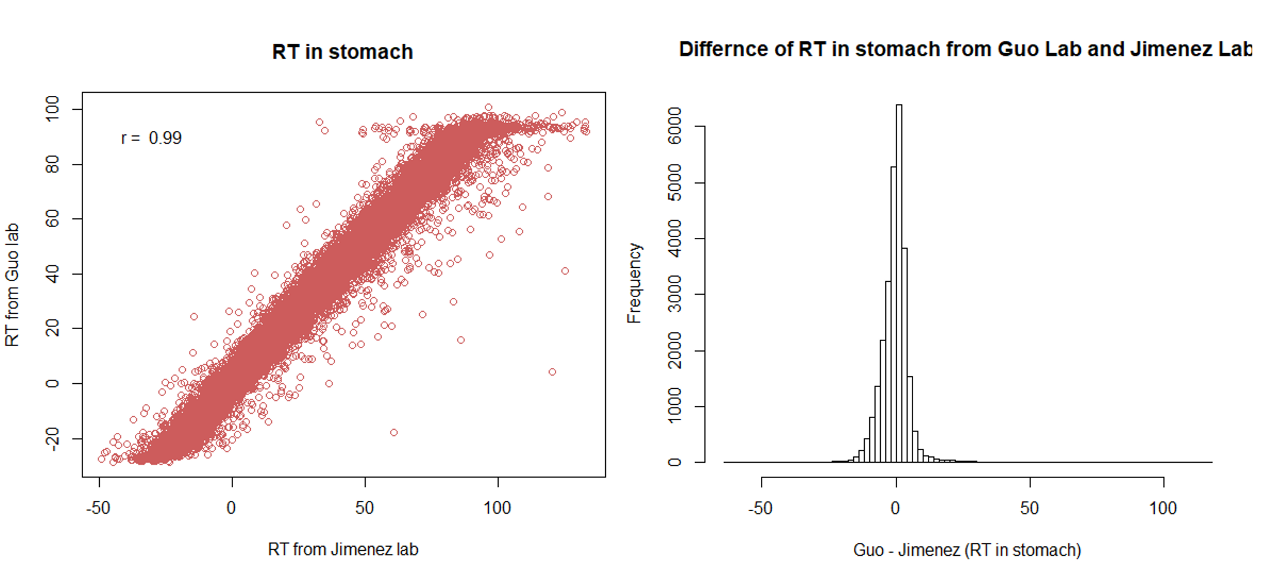

**Figure N2C. Comparison of calibrated RT values from stomach DDA runs acquired in the Guo lab and the Jimenez lab.** Left panel: Peptide precursors identified in both the Guo lab and the Jimenez lab. Right panel: distribution of the Guo/Jimenez ratio of peptide precursors.

Then we checked why the overlap of peptide is not very high. By random selection, we focused on one protein Q14103 with 355 amino acids, and 45.5% sequence overlapped. As shown in Figure N2D, 15 peptides were identified in both labs while 11 and 7 were identified only in the Jimenez and the Guo labs, respectively.

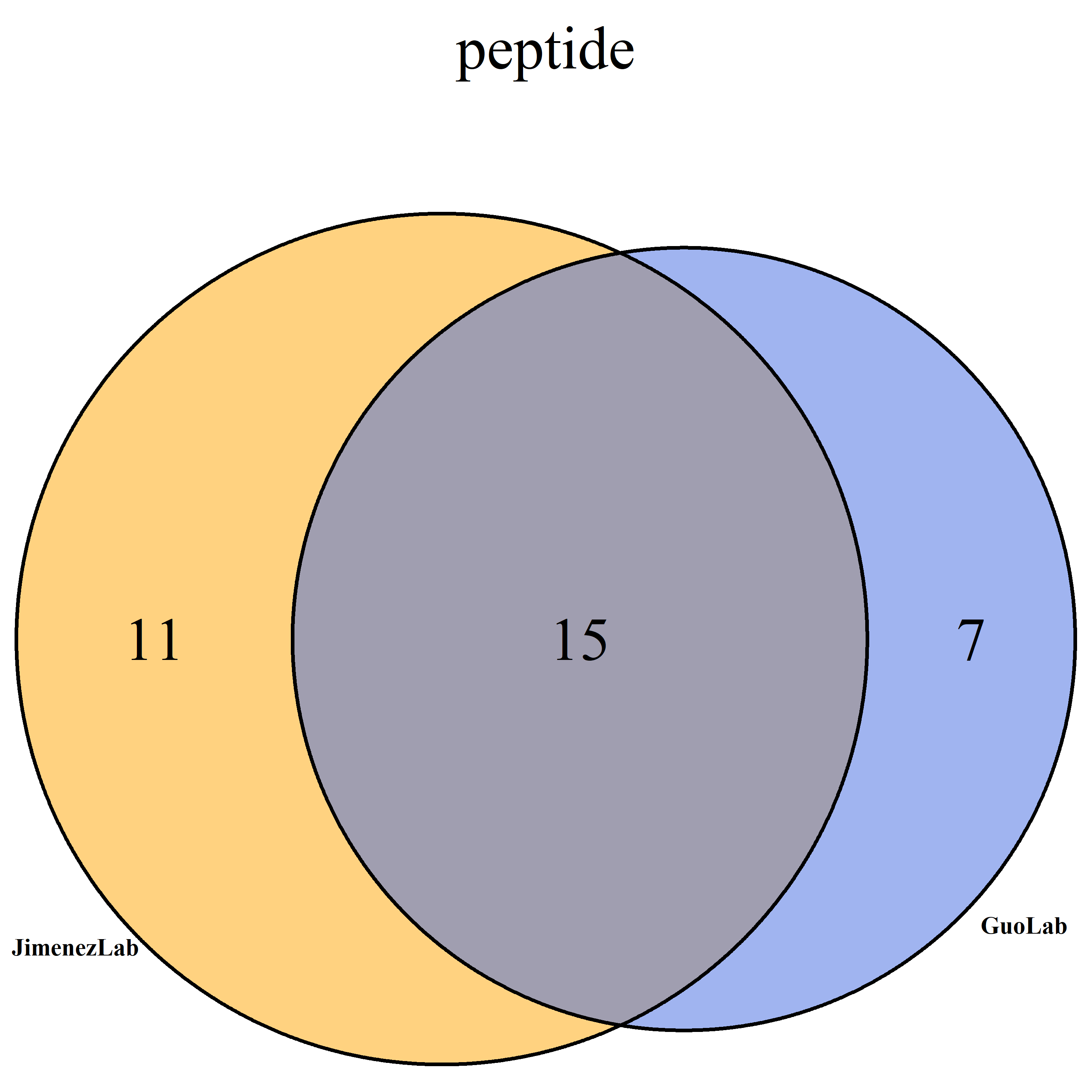

**Figure N2D. Venn diagram of peptide identified from the protein Q14103 in the two labs.**

Then we checked the sequence coverage and found that most of the discrepancy was due to missed cleavage (Figure N2E).

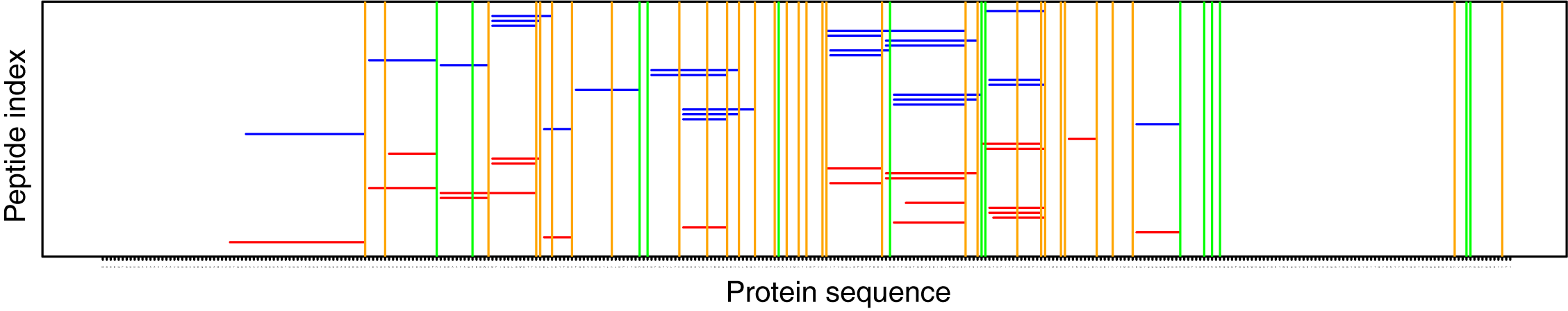

**Figure N2E. Sequence coverage of protein Q14103**. X-axis depicts the protein sequence of Q14103, while the y-axis is the peptide index. Blue horizontal lines represent peptides identified by the Guo lab, while the red lines depict the peptides by the Jimenez lab. Green and orange vertical lines indicate missed cleavage at K and R respectively.

We further investigated the protein sequence coverage in all overlapped proteins. As shown in Figure N2F, proteins identified with higher sequence coverage exhibited higher overlap ratio which is computed using the sequence coverage percentage of overlapped proteins divided by the covered sequence (Pearson correlation coefficient = 0.84).

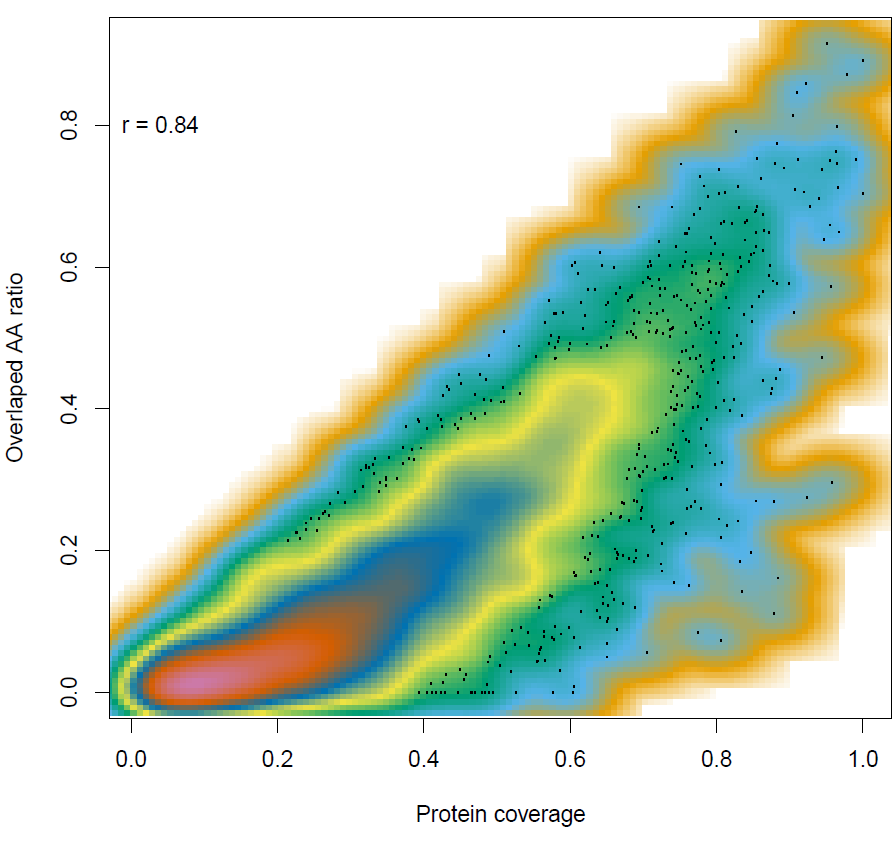

**Figure N2F. The correlation of shared protein coverage with the identified protein coverage in two labs.** The x-axis shows the sequence coverage of proteins identified by the DDA data in the two labs; the y-axis shows the number of residues identified by both labs divided by the protein length. Pearson correlation coefficient was computed.

**Supplementary Note 2. Computational workflow for building DIA library**

**Part 1: Spectral library generation**

1) Obtain the DDA raw data for each tissue type.
2) Retrieve the fasta database. *e.g.* *swissprot_human_20180209_target_IRT_contaminant.fasta*
3) For every tissue type do the following step 4 to 6.
4) Protein and peptide identification using pFind (version 3.1.3)
   a) Parameters for pFind:
      i. Place of decimal: m/z=3
      ii. Uncheck mixture spectra
      iii. Fixed modification= Carbamidomethyl[C]
      iv. variable modification= Oxidation[M]
      v. enzyme=Trypsin_P KR P C
      vi. fasta database as above showed
   b) Obtain the *pFind.spectra* file after pFind is completed.
   c) Convert the raw data to mzXML data using ProteoWizard by running the following command:

*msconvert --mzXML --filter "peakPicking true [1,2] pathToRawData/*.raw*

5) Build the library
   a) Copy pFind.spectra and all mzXML files to the DPHL docker
   b) Run the following command:

*build_library -i pFind.spactra -t iRT.txt*

**Part 2: Build up library with CiRT**

For MS data files without SiRT (iRT), one need to generate CiRT as follows.

1) Build the library as the above steps using the samples with iRT spike-in.
2) Run MaxQuant of representative samples, and obtain the result file *peptides.txt*.
3) Copy the *peptides.txt* and the csv/tsv format of the library built in Part 1.
4) Run the following command in the Docker:

*generate_CiRT dlbcl_peptides.txt dlbcl_library.csv dlbcl*

NOTE: “dlbcl” in this command can be replaced by other tissue type of interest.

5) Then the CiRT library *dlbcl_CiRT.txt* (tsv) will generated.
6) Replace the iRT.txt in the step 5 in Part 1 with this *dlbcl_CiRT.txt* file, and run the step 5 in Part 1.

**Part 3: Build the final library**

When all the sub-library for each sample type are generated, build the consensus library by run the following command:

*buildFinalLibrary.py -i <inputfiles splibs, blank separated and all in one quotes > -w <DIA windows file name.default "/swath/mnt/windows_QE_HF.txt">*

*-o <outputfile name,Optional,default Finallibrary> -c <number of threads, default 4> -h <help>*

For example:

*buildFinalLibrary.py -i dlbcl.splib stomach.splib -w /swath/mnt/windows_QE_HF.txt*

*-o final_consensus*
